# Leveraging Crosslinker Diffusion to Template Stiffness Gradients in Alginate Hydrogels

**DOI:** 10.1101/2024.06.26.599742

**Authors:** Zoe Ostrowski, Tyler Price, Juntao Zhang, Azarnoosh Foroozandehfar, Fred R. Namanda, Tim Kaufmann, Natalia Judka, Tyler Gardner, Mary Thatcher, Emmaline Miller, Lily Mesyk, Abigail Koep, Adam T. Melvin, Juan Ren, Ian C. Schneider

**Affiliations:** Department of Chemical and Biological Engineering, Iowa State University; Beckman Institute for Advanced Science and Technology and Department of Materials Science and Engineering, University of Illinois Urbana-Champaign; Department of Mechanical Engineering, Iowa State University; Molecular, Cellular and Developmental Biology Program, Iowa State University; Cain Department of Chemical Engineering, Louisiana State University; Department of Biochemistry, Bethel University; Department of Chemical and Biomolecular Engineering, Clemson University; Department of Genetics, Development and Cell Biology, Iowa State University

**Keywords:** Stencil, craft cutter, alginic, opacity, durotaxis, mechanotaxis, cancer invasion

## Abstract

Mechanobiology drives many important cell biological behaviors such as stem cell differentiation, cancer drug resistance and cell migration up stiffness gradients, a process called durotaxis. The development of 3D hydrogel systems with tunable 2D mechanical gradient patterns affords the ability to study these mechanosensitive cell behaviors to understand cancer invasion or enhance wound healing through directed migration. In this paper, we developed an approach to spatially imprint within alginate hydrogels, gradients in mechanical properties that can be used to probe mechanobiology. Stencils were easily designed and fabricated using a common craft cutter to control the presentation of a calcium crosslinking solution to alginate gels. Different stencil shapes result in different gradients in opacity that can be imprinted into both thick and thin alginate gels of arbitrary 2D shape. The steepness of the opacity gradient as well as the maximum opacity can be controlled based on reproducible crosslinking kinetics regulated through calcium concentration and gradient developing time. Calcium crosslinking results in both opacity changes as well as increases in elastic modulus in the bulk hydrogel. Opacity correlates with elastic modulus, allowing it to be used as a proxy for local elastic modulus. Functionalized alginate gels with collagen and imprinting stiffness gradients within them resulted in cell invasion that was spatially dependent, where stiffer regions facilitated deeper invasion of breast cancer cells. Consequently, this stenciling approach represents a facile way to control stiffness gradients in alginate gels in order to study mechanosensitive cellular behavior.

**Graphical Abstract:** 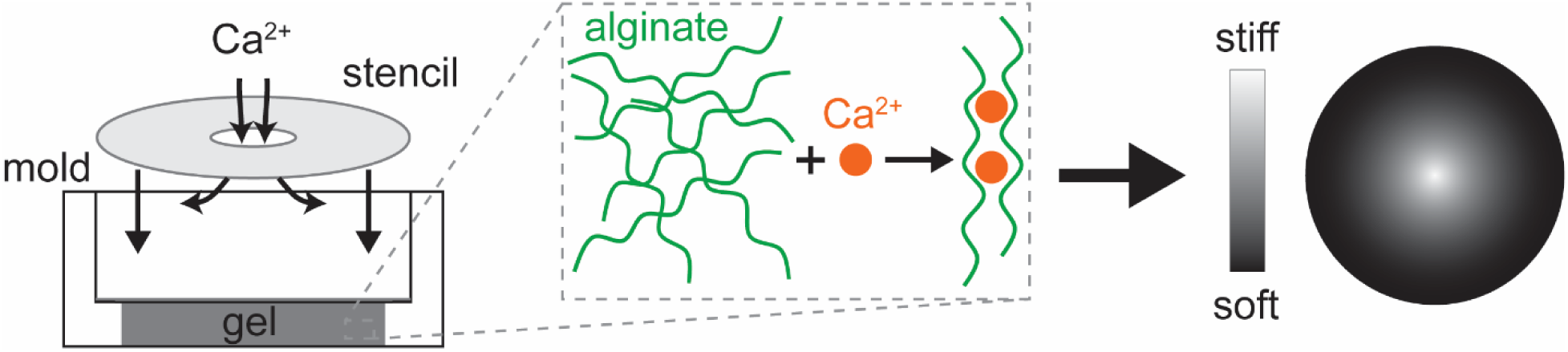

## 1. Introduction

It is well-established that a wide variety of biological processes are driven by mechanobiology. These processes include development[1], immune response[2], wound healing[3] as well as diseases like cancer[4] and fibrosis[5]. Cells sense the stiffness of their environment, altering cell differentiation, drug sensitivity, metabolism and cell migration. Spatial gradients in stiffness formed in biomaterials allow for the study of these cell behaviors. Spatial gradients can reveal non-linear relationships between stiffness and the above mentioned cell behaviors along with the transcriptional, signaling and epigenetic signatures in ways that would be difficult to observe in individual experiments[6–9]. Furthermore, the ability to control gradients in stiffness is absolutely critical for understanding and controlling cell migration through durotaxis[1,10–13], directed cell migration in response to stiffness gradients. Durotaxis drives cancer metastasis as well as stromal cell invasion of the tumor[11]. Furthermore, durotaxis drives wound healing, where stiffness gradients in the wound bed are present and can recruit fibroblasts, organize wound contraction and facilitate hair follicle development[14]. Encoding these types of mechanical property gradients within biomaterials that are provisionally placed in the wound bed will likely lead to enhanced migration and faster wound healing, particularly in traumatic or chronic wounds.

A variety of biomaterials have been used as wound dressings, but the focus of these materials has been on absorption of exudate, maintenance of wound moisture or inhibition of microbial growth[15]. Alginate hydrogels are commonly used in wound dressings[16]. Alginate is a naturally occurring polymer derived from brown algae. Due to its biocompatibility, low toxicity, and structure similarity to extracellular matrices, it is becoming more commonly used in hydrogels for advanced wound dressings[17]. The stiffness of alginate hydrogels can be tuned by crosslinking them with divalent ions like calcium, allowing one to match hydrogel stiffness with tissue stiffness[18,19]. However sophisticated hydrogels with arbitrary stiffness gradients that could direct cells to particular targets within the wound bed at a faster rate have not been developed. In order to study the cellular and molecular mechanisms of cell migration that are present during wound healing or to generate tissue engineering constructs that can direct cells at high speed to targets, approaches for controlling gradients within hydrogels will need to be developed.

There are many ways to form biological gradients[20]. The most commonly used hydrogel system in which to generate mechanical property gradients and study mechanobiology or durotaxis is polyacrylamide. Stiffness gradients can be generated easily using designed masks and photoinitiators[21]. However, while polyacrylamide is a good model hydrogel system, cells cannot be readily embedded within it due to toxicity issues during polymerization and cell invasion is stunted due to an inability for cells to remodel this hydrogel, making its use difficult *in vivo*. Another common system in which to generate stiffness gradients is PEG-polymer systems[6,9], however these can be expensive due to the cost of functionalized PEG. Stiffness gradients in collagen can be generated in collagen gels as well, however, these spatial gradients are usually accompanied by fiber alignment gradients[22]. Alginate hydrogels overcome these problems as they are relatively simple and inexpensive to fabrication. There are several examples of approaches that can generate alginate gradients. Gradients in stiffness have been generated through attachment to stiff interfaces[18,23], using dipping methods[24] or by merging two different alginate solutions, creating a gradient between the two[25]. However, these approaches are limited in the shape of gradients that can be generated. Bioprinting has been used to make gradients in alginate[26,27]. However, while bioprinting is becoming much more accessible for labs, it still requires relatively expensive equipment and optimization. Photopolymerization of alginate can be performed using methacrylated alginate and photoinitiators[28] or photocrosslinking of alginate using embedded vesicles that can release calcium[29]. However, while these approaches afforded very good spatial control and even allowed for temporal control, they either used reagents that can be cytotoxic or difficult to produce. What is needed in order to form stiffness gradients in hydrogels to study mechanobiological responses of cells is a system that can generate a wide variety of gradients, does not require expensive equipment and is easy to use.

Here we develop a stenciling method that can generate unique and tunable stiffness gradients within alginate hydrogels leveraging the controlled diffusion of calcium into the gel. The stencils are easily designed and fabricated, producing a variety of patterns. We show that calcium cross-links the alginate, resulting in an opacity change due to local polymer contraction or densification and enhanced light scattering. We have generated different gradient patterns in both thick and thin systems and characterized the kinetics of crosslinking. Bulk mechanical property measurements have been made and we have spatially mapped mechanical properties to the spatial position and have correlated this to changes in opacity. Finally, to we have demonstrated a spatially controlled cell biological response, cancer cell invasion, demonstrating the utility of forming these gradients in mechanical properties. This technique allows for the easy design and fabrication of alginate gels with tunable stiffness gradients.

## 2. Materials and Methods

### 2.1 Assembly of stenciled alginate hydrogels

A custom 1-inch diameter mold with a 15.4-mm diameter hole was filled with 1 ml of an alginate gel containing 20 mg/ml alginate (alginic acid sodium salt from brown algae, Sigma Aldrich, low viscosity, MW ∼ 30-100 kDa, A1112) and 20 mM calcium sulfate (CaSO_4_, Acros Organics, 385351000). Additional buffers were used to dissolve the alginate and CaSO_4_ including phosphate buffer saline (PBS) lacking magnesium and calcium (Gibco, 14190144) and Dulbecco’s Modified Eagle’s medium (DMEM, Sigma Aldrich, D6546) with 1% penicillin-streptomycin (Gibco, 15140122) and 10% heat-inactivated fetal bovine serum (FBS, Gibco, A5256801). A custom cut stencil is placed on top of the gel and sealed to the edge of the mold with vacuum grease (Dow Corning). Stencils are cut on transparency paper (Apollo Write on Transparency Film Clear) using a Cricut Maker 3. The Cricut Maker 3 allows for creation or uploading of any custom designs, shapes and text and has a spatial resolution of 1 mm diameter for the circle design (**Supplementary** Figure 1). All stencils have an overall diameter of 1 inch to fit the diameter of the mold. The smaller circle stencil has a diameter of 2.5 mm. The larger circle stencil has a diameter of 5 mm. The line stencil has a width of 1 mm, and the S shape stencil has a width of 3 mm. On top of the stencil is a solution of 80 mM CaCl_2_ (CaCl_2_ Fisher Scientific, C79) (**Figure 1A**). Thin alginate gels were generated using a mold cut using the Cricut Maker 3. Two layers of 1 mm thick silicone attached by two-sided adhesive were stacked together to create a chamber for a thin alginate gel. Thin alginate gels were made with final concentrations of 20 mg/ml alginate and 20 mM CaSO_4_. A stencil made from transparent cellulose acetate was placed on top of the alginate gel, and 80 mM CaCl_2_ filled the 35 mm cell culture dish (**Figure 2A**). Images were taken every five minutes for 8 h.

**Figure 1:**
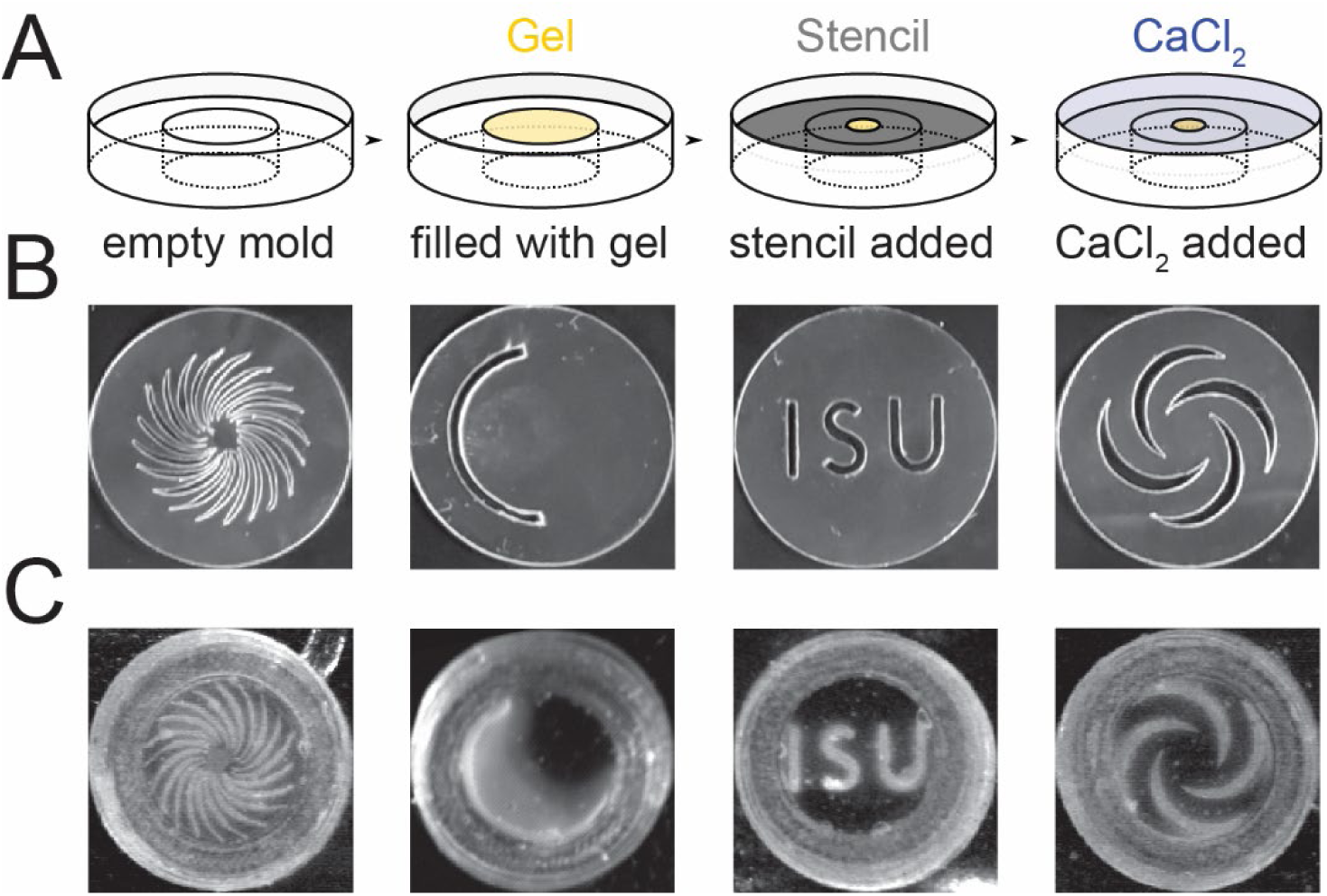
Approach for stenciling opacity gradients into alginate gels. A) Schematic of approach to fabricate alginate gels. Alginate gels (2700 - 5400 μm thick) are initially sparsely cross-linked using low calcium concentration (20 mM CaSO_4_) in molds (yellow). A stencil is placed above with different geometries of openings (grey). A solution of high calcium concentration (80 mM CaCl_2_) is placed on top of the stencil to spatially cross-link the gel (blue). B) Images of uniquely designed stencils made of transparency film (cellulose acetate, 10 μm thick) cut using a craft cutter. C) Images of alginate hydrogels spatially cross-linked using stencils in B) after > 1h 20 min of incubation. Images are 25.4 mm by 25.4 mm.

**Figure 2:**
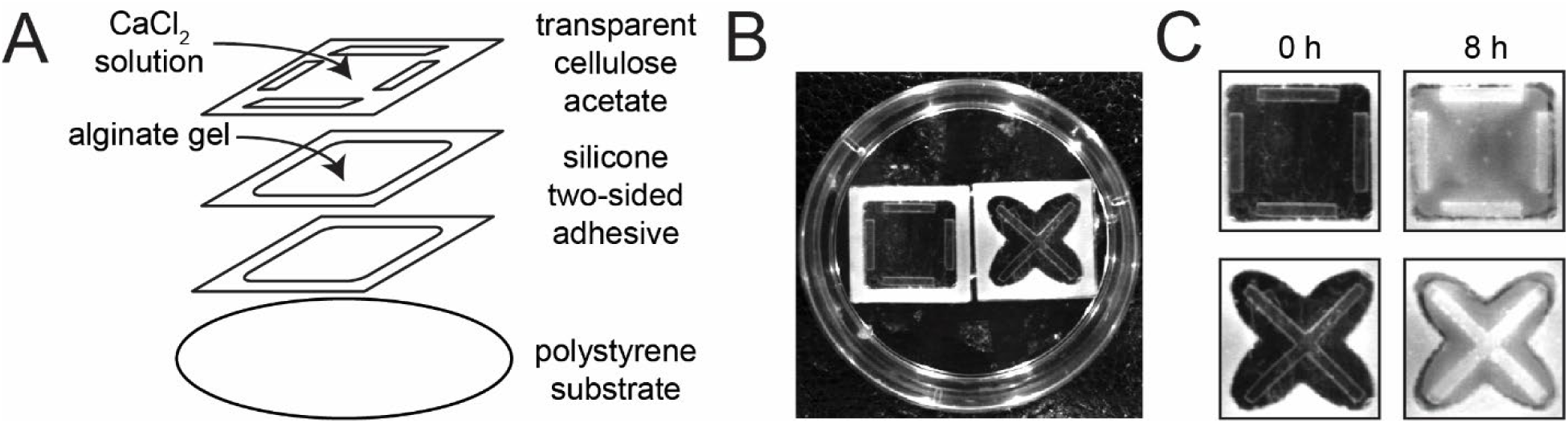
Stencils can generate opacity gradients in thin alginate gels. A) Two–sided adhesive silicone sheets cut with patterns were doubly stacked creating a 2 mm thick reservoir to hold sparsely cross-linked alginate gels (20 mg/ml alginate gelled with 20 mM CaSO_4_). This reservoir was adhered to a polystyrene cell culture dish. A stencil was adhered to the top silicone sheet and the cell culture dish was filled with 80 mM CaCl_2_ to crosslink alginate gel exposed to the solution. B) Image of two chambers attached to one 35 mm polystyrene cell culture dish. The silicone well is white. C) Images were taken before crosslinking (0 h) and after crosslinking (8 h) with 80 mM CaCl_2_.

### 2.2 Fabrication of 3D-printed imaging chambers

A 3D printed imaging chamber was fabricated and adhered directly to a glass slide. The 3D printed imaging chambers were designed in SOLIDWORKS CAD software (3DS, Paris, France) and printed using Overture PLA (polylactic acid) 1.75 mm diameter filament with the Creality Ender-3 V2 3D printer. The .stl files from SOLIDWORKS were converted to .gcodes formed for printing in the slicing software Simplify3D (Simplify3D, Cincinnati, OH, USA). Two varieties of imaging chamber were fabricated: thin and thick. Both thin and thick chambers were printed with an imaging aperture dimension of 18 mm in diameter, an interior shelf width of 3 mm, a wall thickness of 1 mm, for a total diameter of 28 mm. Thin chambers had a total wall height of 5 mm, including the interior shelf thickness of 2 mm whereas thick chambers had a total wall height of 8 mm, including the interior shelf thickness of 4 mm. After printing the imaging chambers were removed from the printer bed and adhered to a 75 mm x 25 mm x 1 mm glass slide using vacuum grease. This forms the completed imaging chamber-slide complex that allows for the visualization using light microscopy.

### 2.3 Imaging and image analysis of alginate gels

Alginate gels were imaged using a home-built system including an 8-bit video camera (Javelin CCD) over a period of 8 h with images taken every 5 min. The initial image of the gel was taken immediately before 80 mM CaCl_2_ was placed on top of the stencil. Using ImageJ, the change in opacity (intensity) was measured as a change in gray value at three locations on each gel: the center of the stencil, 1 mm from the edge of the stencil and 3 mm from the edge of the stencil. The intensity was normalized by the following equation:

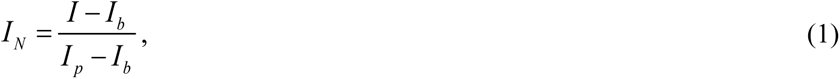

where *I_N_* is the normalized brightness, *I* is the brightness at a particular position and time, *I_b_* is the background intensity, and *I_p_* is the peak intensity. For **Figure 3**, *I_b_* is the intensity at time equal 0 in the center of the stencil and *I_p_* is the peak intensity at time equal 8 h in the center of the stencil. For alginate gels with no stencil and no CaCl_2_, *I_p_* values were from images taken of alginate gels with no stencil and 80 mM CaCl_2_. For **Figure 3**, *I_b_* is the average over the first hour between 0 and 4 mm away from the center and *I_p_* is the average intensity between 12 h and 13 h between 0 mm and 1 mm. The absolute gradient is the slope of the normalized intensity as a function of radial distance (**Supplementary** Figure 4). Degradation of alginate gels was imaged the same way as described above. After 8 h of 80 mM CaCl_2_ crosslinking, the solution and 5 mm diameter circle stencil were removed (*t* = 0 h) and the gels were placed in either water, PBS, or 10% DMEM and imaged every 5 min for 8 h. Using ImageJ, the change in opacity (intensity) was measured as described above at the same three locations. The intensity was normalized by the following equation:

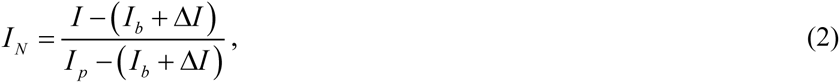

where *ΔI* is the change in intensity due to the change in buffer added at *t* = 0 when either water, PBS or 10% DMEM is added (**Supplemental Figure 5**).

**Figure 3:**
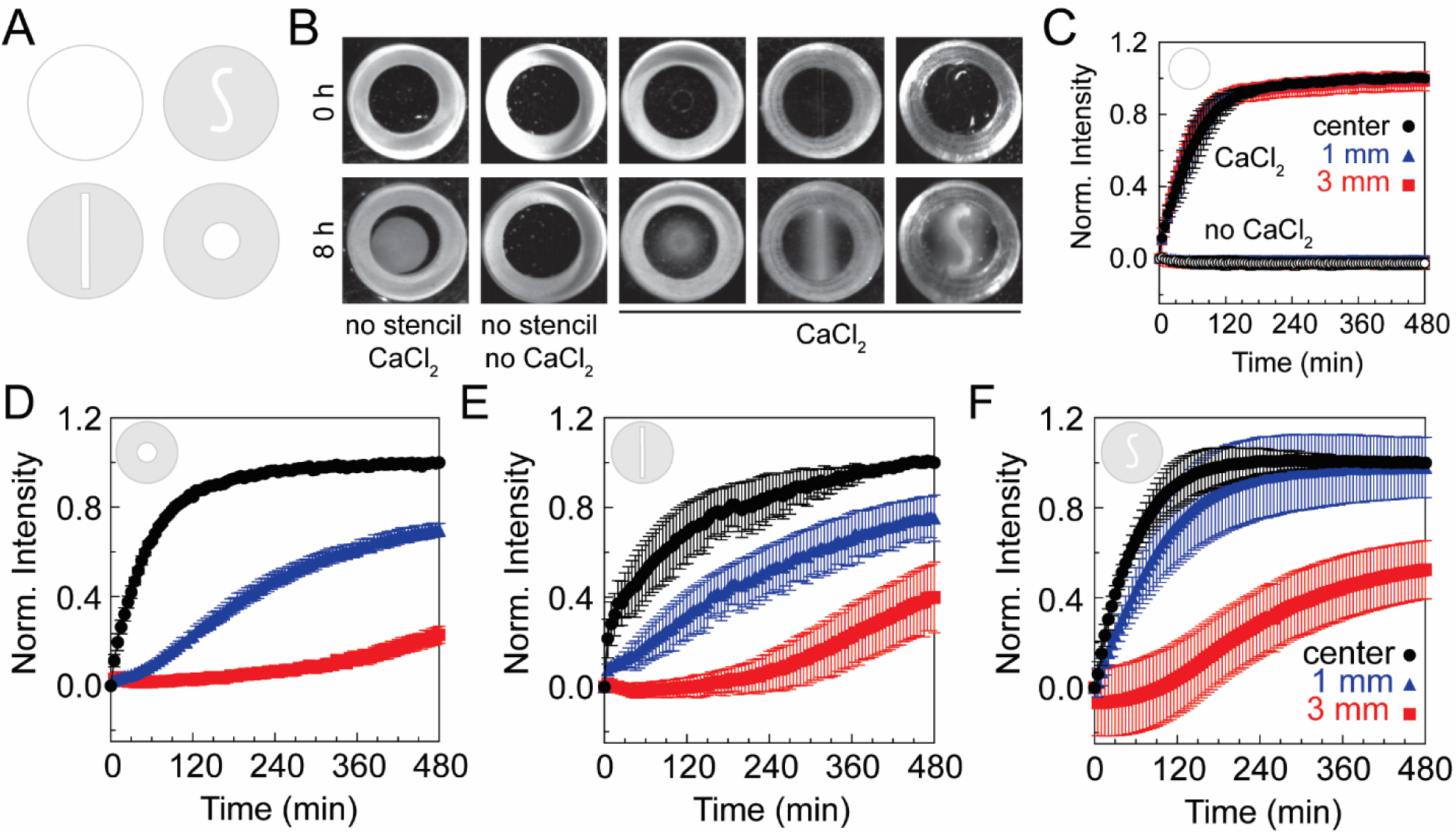
Stencil shape and exposure time determine the opacity gradient in alginate gels. A) Schematic of the unique stencils used to create gradients in opacity within the alginate gels. B) Representative images of alginate gels (20 mg/ml alginate cross-linked with 20 mM CaSO_4_) before (0 h) and after (8 h) exposure to an 80 mM CaCl_2_ solution above the stencil. Images are 25.4 mm by 25.4 mm. C) Normalized intensity over time with no stencil on the alginate gels and with and without 80 mM CaCl_2_ at the center (black circles), 1 mm (blue triangles) and 3 mm (red squares) from the edge of the stencil (*N_days_* = 3, *N_gels_* = 3, *N_points, center_* = 3 and *N_points, 1 mm or 3 mm_* = 6). Normalized intensity over time for the D) circle stencil (*N_days_* = 2, *N_gels_* = 5, *N_points, center_* = 5 and *N_points, 1 mm or 3 mm_* = 10), E) line stencil (*N_days_* = 5, *N_gels_* = 5, *N_points, center_* = 5 and *N_points, 1 mm or 3 mm_* = 10) and F) S-shaped stencil (*N_days_* = 5, *N_gels_* = 5, *N_points, center_* = 5 and *N_points, 1 mm or 3 mm_* = 10) at distances 0 (center), 1, 3 mm from the center of the gel. Error bars represent 95% confidence intervals.

### 2.4 Measurement of the absorbance of alginate gels during polymerization

The kinetics and extent of crosslinking alginate with CaSO_4_ and CaCl_2_ were analyzed using a UV-Vis Spectrophotometer (Varian Cary 50 Bio) and polystyrene disposable cuvettes (Semi-Micro, 1 cm path, 1.5 ml, 12.5 mm square x 45 mm high; Bio-Rad, 2239955). The alginate and CaSO_4_ were mixed, and the resulting 1 ml gel in the cuvette had a final concentration of 20 mg/ml alginate and 20 mM CaSO_4_. A 1 ml solution of 180 mM CaCl_2_ or 80 mM CaCl_2_ filled the top of the cuvette. The scans were started immediately following the mixing of the alginate and CaSO_4_ (**Supplementary** Figure 2).

### 2.5 Elastic modulus measurement in bulk alginate gels

Uniaxial compression testing was used to measure elastic modulus in bulk alginate gels. The mold was removed from around the gel and a coverslip was placed on top. Compressive force was applied by placing individual weights on top of the coverslip until deformation was observable. The weight was incrementally increased until failure. The alginate gels were imaged 60 frames per second. (**Figure 5A**) Assuming a circular cross section, the engineering stress (*σ*) in the gel was calculated using the following equation:

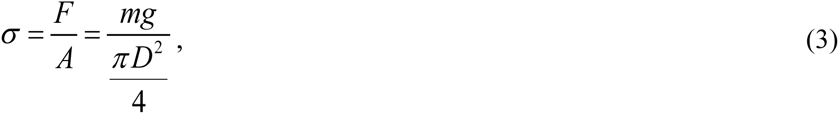

where *m* is the mass of weight, *g* is acceleration due to gravity, *A* is the initial gel area and *D* is the diameter of the top surface of the undeformed gel measured from the side view. The engineering strain (*γ*) was calculated using the following equation:

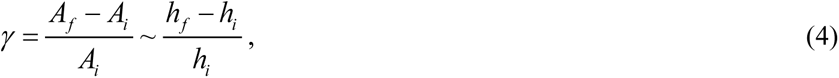

where *A_f_* and *A_i_* are the final and initial projected trapezoid area from the side and *h_f_* and *h_i_* are the average heights of the final and initial trapezoid shape (**Figure 5B**). Elastic modulus was calculated using the following equation:

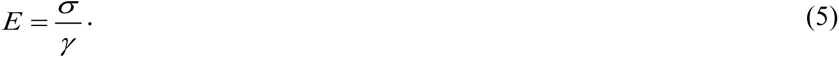

### 2.6 Functionalization of coverslips covalent attachment of alginate to coverslips

Alginate gels float in both water and NaCl solutions. To prevent any movement during mechanical property measurements, the alginate gels were adhered to functionalized glass coverslips[30]. First, glass coverslips were submerged in piranha solution composed of 3:1 sulfuric acid (H_2_SO_4_, Fisher Scientific, A300) and 30 wt% hydrogen peroxide (H_2_O_2_, Fisher Scientific, H325) for 1 h. Then the coverslips were rinsed with deionized water three times before placing a solution of 1% aminoproplytriethoxysilane (APTES, Acros Organics, 43094) in 0.174 M acetic acid (Fisher Scientific, A38) on top of the coverslips for two hours. Afterwards the coverslips were rinsed in deionized water three times. Second, alginate was attached to functionalized glass coverslips in the following way. A 1 wt% alginate solution was prepared in 0.1 M 2-(N-morpholino)ethanesulfonic acid buffer (MES, Sigma Aldrich, M8250 ). Sulfonated N-hydroxysuccinimide (NHS, Sigma Aldrich, 130672) and 1-ethyl-3-(3-dimethylaminopropyl) carbodiimide (EDC, Thermo Scientific, 22980) were added to the alginate solution in a molar ratio of 1 alginate:30 NHS:25 EDC, and brought to a pH of 6.5 using 1 M NaOH (Fisher Scientific). To assemble the functionalized alginate gels, a coverslip was placed in the bottom of the mold and 75-100 µL of the functionalized alginate solution was placed on top of the functionalized coverslip so that there was a thin layer of solution covering the entire surface area of the coverslip. The alginate gels were prepared as described above and the initial gel was placed on top of the functionalized alginate on the coverslip.

### 2.7 Characterization of functional attachment of alginate to glass

Attenuated Total Reflectance-Fourier Transform InfraRed spectroscopy (ATR-FTIR) was performed on an FTIR spectrometer (Nicolet MAGNA 750, Thermo Scientific) and attached variable-angle specular reflectance accessory (Pike VeeMax III, Pike Technologies, Madison, WI, USA). ATR-FTIR was used to assess whether dry, wet or unfunctionalized glass coverslips contained surface amines. In addition, the density of amines on the surface of the coverslips was quantified using UV absorbance of 4-nitrobenzaldehyde (4-NB, Sigma Aldrich) attached to amines on functionalized coverslips as we have reported elsewhere[31]. Calibration curves of absorbance of 4-NB were generated reading spectra from a UV-Vis Spectrophotometer (Varian Cary 50 Bio). To assess the degree of amine functionalization, functionalized glass coverslips were immersed in a solution of 25 ml of anhydrous ethanol containing 20 µl of acetic acid and 0.4 mg/ml 4-nitrobenzaldehyde (4NB, Sigma Aldrich) at 50 °C for 3 h. The coverslips were then washed with absolute ethanol and dried with air and crushed and placed in a solution of 0.2% acetic acid for 1 h at 30 °C. The absorbance of 4NB was measured and the calibration curve was used to calculate the number density of 4NB and consequently amines on the surface (**Supplementary** Figure 6)[32].

### 2.8 Measuring swelling of alginate gels

Due to a difference in osmotic pressure, alginate gels swell in water. To minimize the effects of swelling on the mechanical property measurements, the alginate gels were submerged in a salt solution. Alginate gels were made with 20 mg/ml alginate and 20 mM CaSO_4_. No stencils or CaCl_2_ was used in these experiments. The weight of the initial gel was measured before a sodium chloride (NaCl, Fisher Scientific, S271) solution was added for 12 h. Then the salt solution was removed, and the final weight of the gel was measured. The percentage swelled is calculated by dividing the mass of the water absorbed by the initial mass of the gel (**Supplementary** Figure 7).

### 2.9 Local elastic modulus measurements in alginate gels

Spatial mechanical properties were measured using Atomic Force Microscopy (AFM). Functionalized alginate gels were prepared as described above, and 80 mM CaCl_2_ was placed on top of the stencil for 3 hours. Images were taken before and after adding and removing the 80 mM CaCl_2_. The brightness measurements from the before and after images were analyzed as described above. The gels were placed into a 60 mM NaCl solution to minimize swelling for the duration of AFM testing. The force measurements were performed in contact mode on a commercial AFM platform (Bio-Resolve, Bruker Nano, Inc.). The AFM cantilever was 0.03 N/m. The ramp speed (i.e., force loading speed) and ramp rate were 100 nm/s and 0.1 Hz, respectively. To obtain elastic modulus, *E*, the experimental deflection, and height data was modeled by:

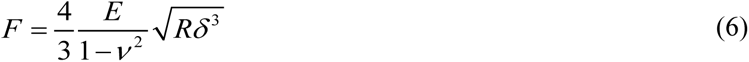

where *F* is the force, *E* is the Young’s modulus (elastic modulus), *ν* is Poisson’s ratio (0.45[33]), *R* is the radius of spherical tip (5 μm) and *δ* is the indentation depth. Indentation depth is defined as the height of the probe minus the deflection. To calculate the elastic modulus, the force was plotted as a function of adjusted area. Adjusted area is a function of indentation depth given by:

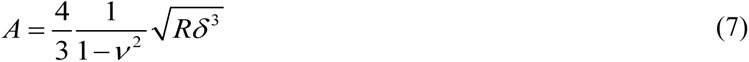

The slope of the force versus adjusted area is the elastic modulus (**Supplementary** Figure 8). Three locations on each gel were measured with AFM. For the circle stencil the locations were the center of the stencil, 1 mm and 3 mm from the edge of the stencil. For the uniform gels without a stencil, the locations were the center of the gel, and 3 mm to the left and right of the center of the gel.

### 2.10 Collagen functionalization of alginate gels

Alginate (40 mg/ml) was functionalized with EDC/NHS as described above. Collagen type I solution (Cellink) was combined with 0.1 M MES buffer and 40 mM CaSO_4_ at a pH of 6.5. Alginate gels using the functionalized alginate solution and CaSO_4_ solutions were made as described above. A 80 mM CaCl_2_ solution was placed on top of the 5 mm diameter circle stencil for three hours at room temperature. Then the stencil was removed, and the gels were pinned in place with a 14 mm diameter cut stencil to prevent floating and movement of the gel since functionalization of glass as described above fails under prolonged incubation at 37 °C. The gels were incubated in 10% DMEM for 12 h at room temperature before cells and fresh media were added.

### 2.11 Cell culture and staining

MDA-MB-231 cells (human mammary carcinoma cell line, ATCC) were cultured at 37 °C with 5% CO_2_ in DMEM with 10% heat-inactivated fetal bovine serum (FBS) (Gibco), 1% GlutaMAX (Gibco), and 1% penicillin/streptomycin (Gibco). The samples were prepared with cells from subcofluent cultures (70-90%) and plated at a density of 40,000-80,000 cells/cm^2^. Cells were fixed with 4% paraformaldehyde (Thermo Scientific) in a cytoskeleton buffer consisting of 10 mM MES (Fisher, pH 6.1), 3 mM MgCl_2_ (Fisher Scientific), 138 mM KCl (Fisher Scientific), and 2 mM Ethylene Glycol Tetraacetic Acid (EGTA, Sigma Aldrich). Next, cells were permeabilized using a 0.5% Triton-X cytoskeletal buffer for 5 min. For staining, cells were incubated with Alexa 488-phalloidin (Thermo Fisher Scientific, A12379) in 1x Tris Buffered Saline (TBS, 140 mM NaCl (Fisher Scientific) and 15 mM Tris base (Fisher Scientific), pH 7.5) containing 0.1% Tween-20 (Fisher Bioreagents) v/v and 2% bovine serum albumin (Sigma Aldrich,) w/v for 1 h. A circle coverslip with a diameter of 25 mm (Fisher Scientific, 1241010) was used to seal the gel inside the mold. The chamber was flipped upside down for imaging with confocal microscopy.

### 2.12 Cell imaging and analysis

Live cells were imaged using a Nikon Ti-E inverted microscope with a 10x phase contrast objective (*NA* = 0.3). Images of cells on the surface of the gels were taken across the diameter of the collagen functionalized alginate gel. The images were stitched together using ImageJ/Fiji [34] at 4 and 24 h. The total number of cells and their location relative to the edge of the stencil were recorded. After fixing and staining, the cells on collagen functionalized alginate gels were imaged using a Leica Stellaris Stimulated Emission Depletion (STED) confocal microscope with a 10x air objective (*NA* = 0.3). Images were taken at three different locations on the gel: center, 1 mm and 3 mm from the edge of the imprinted gradient. Images were taken at different depths at each location starting at the surface stepping every 10 μm into the gel for a total of 21 images. The number of cells at each depth at 4 and 24 h was normalized to the functionalized collagen gel at 4 or 24 h, respectively using the following equations:

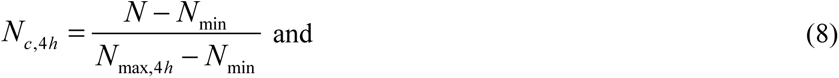

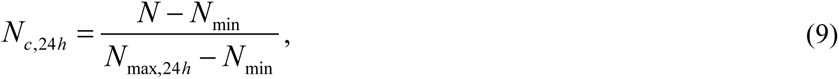

where *N_c_* is the normalized number of cells at either 4 or 24 h, *N* is the number of cells at a particular location and depth, *N_min_* is the number of cells at 4 h, at the location of 3 mm from edge of stencil and at depth of 200 μm inside the gel, *N_max_* is the maximum number of cells at any depth within the gel and at any location on the gel at 4 and 24 h, respectively.

### 2.13 Statistical analysis

The significance level is set to *p* = 0.05. Box plots: Upper and lower error bars are minimum and maximum values, top and bottom of the box represent the 75^th^ and 25^th^ percentile, the line is the median and the dashed line is the mean. The error bars shown on the figures are 95% confidence intervals. *T*-tests were used to gauge statistical significance of mechanical property difference and statistical significance was set at *p* = 0.05 and denoted with an asterisk.

## 3. Results

### 3.1 Unique stencil patterns result in tunable opacity patterns

Gradients in alginate crosslinking were generated leveraging diffusion of calcium through custom designed stencils. Alginate (20 mg/ml) sparsely crosslinked with calcium sulfate (20 mM CaSO_4_) was loaded into molds with a recessed shelf (**Figure 1A**). This shelf allows for the application of a clear stencil on top of the gel. The stencil contains a customizable pattern cut into cellulose acetate using a craft cutter (**Figure 1B**). This particular craft cutter is able to cut sizes as small as 1 mm (**Supplementary** Figure 1). A high concentration calcium chloride solution (80 mM CaCl_2_) was placed on top of the stencil. Calcium diffuses into the gel through the pattern, creating a gradient in opacity in the *x*-*y* plane (**Figure 1C**). The opacity was quantified by calculating the intensity of the alginate gel at a particular position and time. Opacity is highest underneath the pattern and increased with time. In addition to opacity gradient formation in thick alginate gels in molds, opacity gradients also formed in thin alginate gels (2 mm) fabricated in silicone wells cut to different shapes (**Figure 2A**). Thin multilayer chambers with both unique well patterns and unique stencils were assembled in petri dishes. Sparsely crosslinked alginate was loaded into these thin wells, overlaid with a stencil and submerged in high concentration calcium chloride (**Figure 2B**). This resulted in increased intensity over a period of 8 h, producing a gradient in intensity templated by the stencil pattern (**Figure 2C**). This demonstrates that opacity gradients can be imprinted into either thick or thin gels by tuning the stencil pattern and duration of gradient development, allowing for a tremendous amount of flexibility in creating opacity gradients with different shapes and gradient steepness.

### 3.2 Characterization of opacity gradients and kinetics of change in opacity

Given that unique gradients can be formed using different stencil patterns, the kinetics of spatial pattern formation was assessed using both absorbance of alginate gels within cuvettes and intensity of alginate gels imaged within the molds over time. Intensity from images at different positions and at different times was normalized to the maximum intensity within the alginate gel. Alginate gels sparsely crosslinked with two different concentrations of calcium sulfate and exposed to different concentrations of calcium chloride showed that above 50 mM CaCl_2_ a maximum level of absorbance was attained, indicating that the high concentration calcium chloride solution (80 mM CaCl_2_) used above was saturating, so this concentration was selected for further experiments (**Supplementary** Figure 2A). Four different stencil model patterns were used (**Figure 3A**). Sparsely crosslinked alginate gels exposed to high concentrations of calcium chloride without a stencil produced a high intensity alginate gel with no gradient (**Figure 3B**). The contraction of the alginate gel upon crosslinking is revealed by the thin black crescent formed when the gel pulled away from the mold. Alginate gels exposed to water instead of calcium chloride resulted in a gel with no change in intensity within the gel. To characterize the spatial kinetics of intensity changes, the normalized intensity of these gels was calculated at the center and at 1 and 3 mm away from the edge of the stencil between 0 and 8 h (**Figure 3C**). The alginate gels exposed to calcium chloride with no stencil had the same rate of increase in intensity at all three locations. After 2 h, the intensity within the alginate gels reached its maximum and remained constant for the remaining 8 h. Similar kinetics were observed when measuring the absorbance within cuvettes of the sparsely crosslinked alginate gels exposed overtop to high concentrations of calcium chloride (**Supplementary** Figure 2B). Interestingly, two behaviors were observed in the cuvette absorbance experiments, one that matched the kinetics of the imaging in **Figure 3C** and one that was much slower. The slow increase in absorbance is consistent with a moving front of opacity oriented in the downward position of the cuvette, while the fast increase is consistent with a moving front of opacity oriented from the sides of the cuvette. Alginate gels contract after crosslinking, resulting in an occasional pulling away from the sides of the cuvette, creating a channel along the side that filled with high concentration calcium chloride. Indeed, a simple diffusion rate calculation based on the diffusion coefficient of calcium (1300 μm^2^/s) and different length scales (height = 3 mm vs half-width = 1 mm) for the two scenarios predicts a lag of 1-2 h as compared to the fast kinetics. This is precisely what is observed (**Supplementary** Figure 2B). Finally, the alginate gels that were not exposed to CaCl_2_ produce no change in intensity at any of the locations (**Figure 3C**). These data indicate that alginate gels crosslink over 2 h before coming to a steady-state level.

In addition to the no stencil condition, the spatial kinetics of different stencil patterns were quantified. Sparsely crosslinked alginate gels were exposed to high calcium chloride solutions through the circle, line and S stencil (**Figure 3B**) and intensity was measured at the center as well as 1 and 3 mm away from the edge of the stencil (**Figure 3D-F**). The opacity pattern matched the shape of the stencil (**Figure 3B**). In addition to water, alginate gels were assembled in a common buffer (PBS) and cell culture media (DMEM). Intensity gradients formed in these buffers but these gradients were shallower (**Supplementary** Figure 3). For all stencils as the distance from the center of the stencil increases, the increase in intensity over time is slower. The center location, where the gel is directly exposed to the calcium chloride on every stencil increases in intensity with similar kinetics to the no stencil condition. The lag time associated with increases in intensity 3 mm from the stencil edge varied among the stencil shape with the circle stencil being the longest, followed by the line and S stencil (**Figure 3D-F**). The slower kinetics associated with the line stencil as compared to the S stencil is caused by the thinner width of the line stencil compared to the S stencil **(Figure 3E&F**). Quantifying the intensity kinetics for different stencil shapes allow for the selection of a particular time that results in a specific opacity gradient.

Given that the line and S stencil shapes had different widths and resulted in different intensity kinetics far from the center, the effect of the size of the stencil for one particular shape was assessed. Circle stencils with diameters of 5 and 2.5 mm were used to form radial gradients (**Figure 4**). Radial gradients were quantified by averaging linescans over angles between 0 and 2*π* and absolute gradient was quantified by finding the local slope of the radial profile at each point (**Supplementary** Figure 4). The circle stencils showed the characteristic decrease in intensity further from the stencil. Interestingly, these gradients seemed stable, even when the calcium chloride solution was removed and replaced with water (**Supplementary** Figure 5). Large circle stencils showed an increase in intensity far from the stencil for long periods, suggesting the impact of a no flux boundary condition at the outer edge of the mold (**Figure 4A**). This effect was diminished with the smaller circle stencil, indicating that the domain in which diffusion is occurring is essentially semi-infinite for times less than 16 h (**Figure 4B**). Using a simple point source model, the diffusion coefficient in the alginate gel was predicted to be on the order of 160 μm^2^/s, which is about 8-fold lower than that in water, likely slowed by the binding of calcium to alginate. Absolute gradients were calculated for both large and small stencil over time, requiring about 4 h for significant gradient establishment. Gradient steepness increased over time and similar gradients occurred in both small and large circle stencils. Large circle stencils produce robust opacity gradients after 4 h near the stencil edge, but these opacity gradients dramatically flatten far from the stencil. Small circle stencils on the other hand can be treated as semi-infinite point sources that produces robust opacity gradients after 4 h throughout the gel.

**Figure 4:**
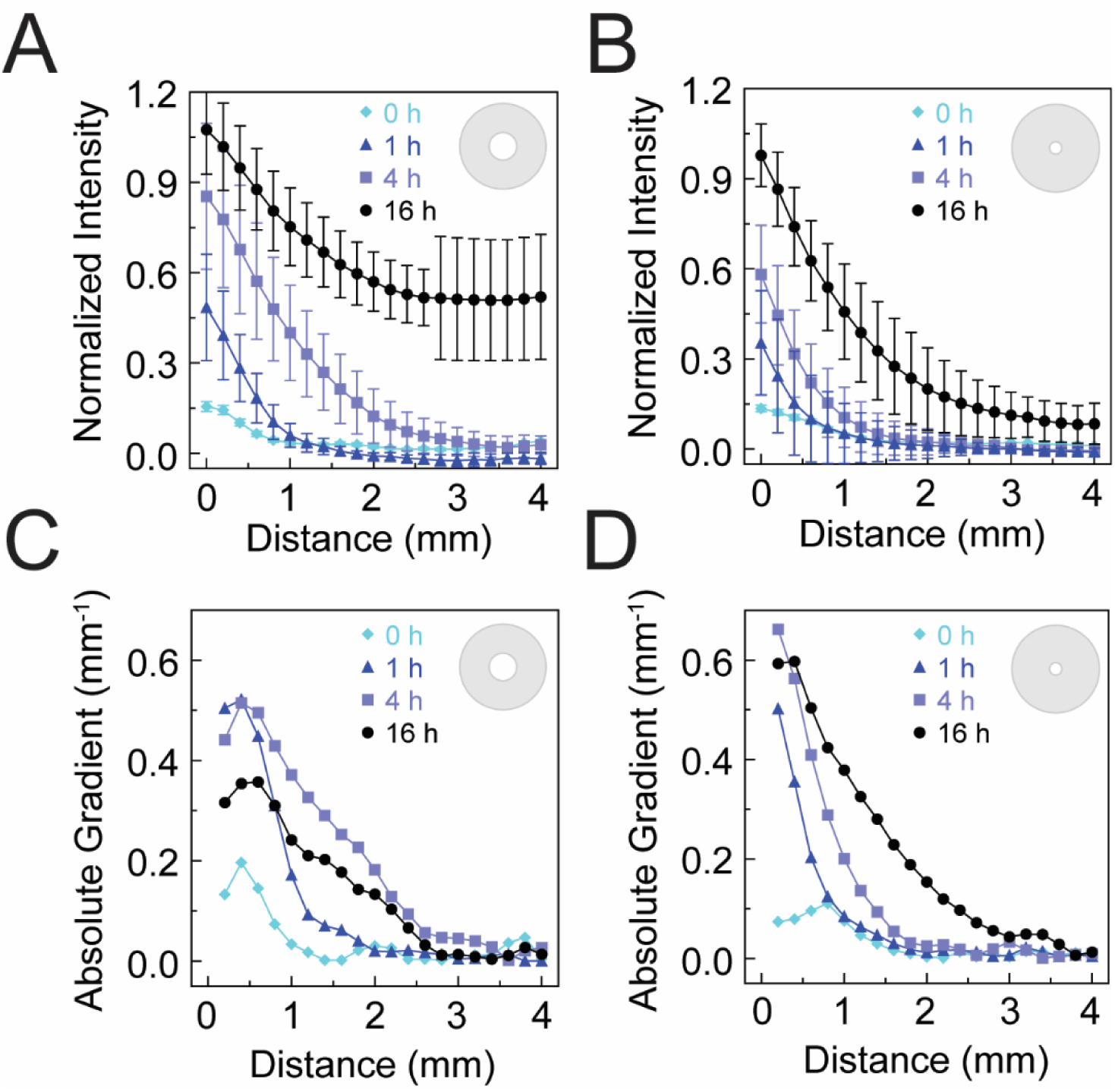
Stencil shape determines the kinetics of opacity gradient development in alginate gels. Normalized intensity within an alginate gel (20 mg/ml alginate cross-linked with 20 mM CaSO_4_) as a function of radial distance from the edge of the opening in a A) 5 mm diameter stencil (*N_gels_* = 8, *N_days_* = 8) and a B) 2.5 mm diameter stencil (*N_gels_* = 8, *N_days_ =* 6) after exposure to an 80 mM CaCl_2_ solution above the stencil. Absolute gradient calculated as the slope of the normalized intensity over radial distance for C) the 5 and D) 2.5 mm diameter circle stencil. Error bars represent 95% confidence intervals.

### 3.3 Characterization of bulk mechanical properties of alginate gels

Given the effects of alginate gel crosslinking on opacity characterized above, we are were interested in measuring the bulk mechanical properties of the alginate gels under similar crosslinking conditions. We used compression testing to assess the compressive elastic modulus of alginate gels. Different masses were placed on top of the alginate gel and the deformation of the alginate was quantified from images (**Figure 5A**). The slope of the stress versus strain curve equals the elastic modulus of the gel (**Figure 5B**). Alginate gels (20 mg/ml) were sparsely crosslinked with different calcium sulfate concentrations (10 mM, 20 mM or 40 mM) in the absence of stencils. The elastic modulus of the alginate gels increased slightly with the concentration of CaSO_4_ (**Figure 5C**). Under conditions used above, the median compressive elastic modulus of the sparsely crosslinked gel is ∼1.5 kPa. When these gels are exposed to high concentrations of calcium chloride, the compressive elastic modulus increased 16-fold (**Figure 5D**). This indicates that alginate gels crosslinked in the stencil system above span roughly 1.5-25 kPa.

**Figure 5:**
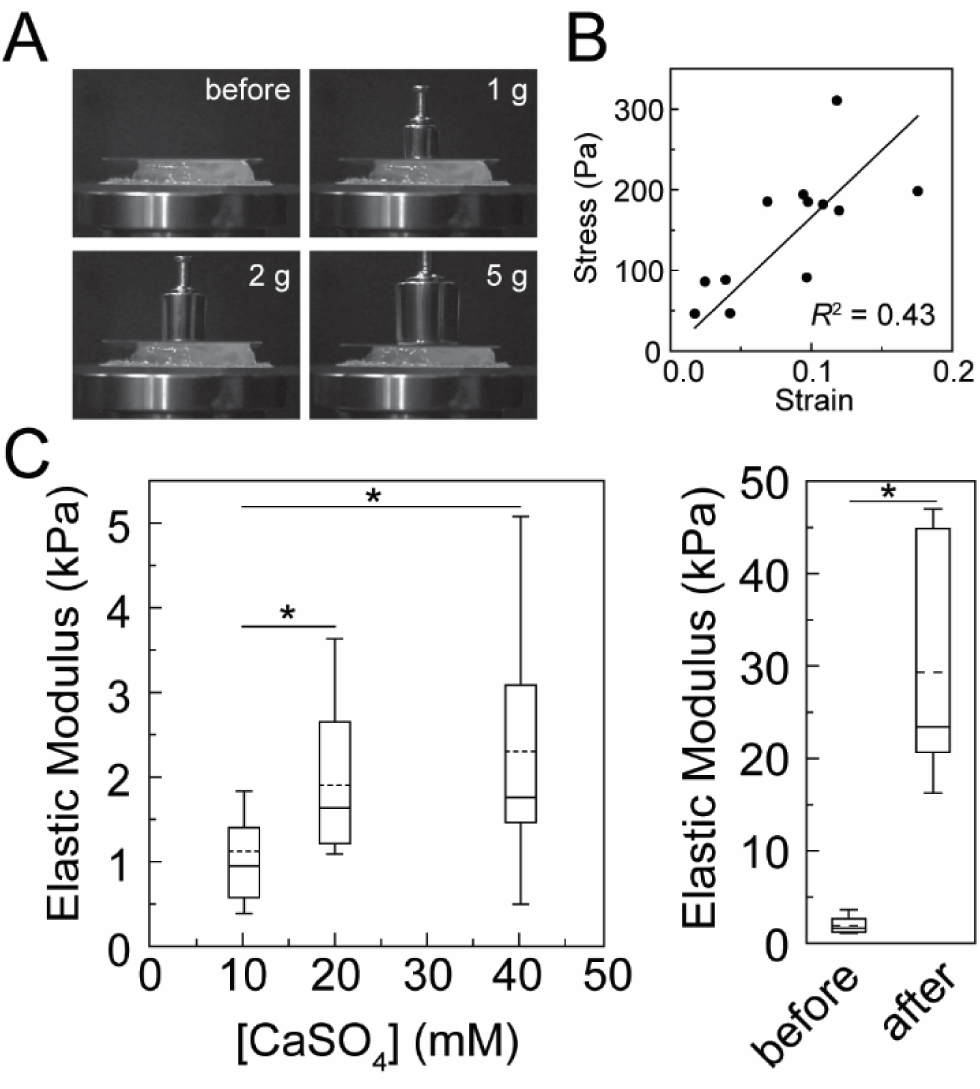
Increasing concentrations of calcium increase elastic modulus of alginate gels. A) Representative images of uniaxial compression testing of alginate hydrogels with different weights. B) Representative stress (*σ*) vs strain (*γ*) curve where the slope is equal to the elastic modulus. Elastic modulus was calculated as follows: *σ* = *Eγ* . C) The elastic modulus of 20 mg/ml alginate hydrogels cross-linked with different concentrations of CaSO_4_ and no CaCl_2_. D) Elastic modulus of 20 mg/ml alginate and 20 mM CaSO_4_ hydrogels before and after (22 h) adding 80 mM CaCl_2_. No stencils were used. *N_gels, 10 mM_* = 14, *N_gels, 20 mM_* = 22, *N_gels, 40 mM_* = 18, *N_gels, 20 mM to 80 mM_* = 7 independent gels. Box plots: Upper and lower error bars are minimum and maximum values, top and bottom of the box represent the 75^th^ and 25^th^ percentile, the line is the median and the dashed line is the mean. Conditions that resulted in *t*-tests with *p* < 0.05 are denoted with asterisks.

### 3.4 Spatial patterns of stiffness can be imprinted into alginate gels

Given the ability to control spatial alginate gel crosslinking using stencils and that this crosslinking results in increases in elastic modulus, we wondered if gradients of elastic modulus were formed and if opacity was predictive of elastic modulus. The elastic modulus was determined at various positions on alginate gels before and after crosslinking with high concentration calcium chloride using AFM. In order to secure the alginate gel to the glass surface, we used EDC/NHS chemistry to attach a layer of alginate to amine functionalized glass. This layer of alginate attaches to the crosslinked alginate gel that was placed over top. This attachment is robust at room temperature, but the attachment is weak if the gel is heated to 37⁰C. We verified glass functionalization by identifying both C-N and –NH_2_ bonds formed after aminosilane chemistry (**Supplementary** Figure 6A). An amine-containing molecule absorbing at around 270 nm (4-nitrobenzyaldeyde (4NB)) was coupled to the functionalized surface, washed, released after hydrolysis and absorbance was measured at 270 nm in order to determine the amine density. Using a calibration curve for 4NB, we determined that the number density of amines to be about 10^7^ #/μm^2^ (**Supplementary** Figure 6B-D). This resulted in alginate gels attached to the glass surface. Indeed, the radial contraction of the alginate gel that is normally observed is not observed when the gel is attached to the glass surface (**Supplementary** Figure 6E). To limit swelling of the alginate gels, gels were submersed in a sodium chloride solution (60 mM NaCl), allowing for AFM measurements to be done in solution (**Supplementary** Figure 7). Using AFM, a spherical probe attached to a cantilever was brought in contact with the alginate gels and the height and deflection of the cantilever was measured (**Supplementary** Figure 8). This allowed for the measurement of elastic modulus in solution at multiple positions.

First, sparsely crosslinked alginate gels were exposed to a high concentration calcium chloride. Images were taken before and after crosslinking (**Figure 6A**). The characteristic increase in opacity after crosslinking was observed. The elastic modulus was calculated at the center of the gel and 3 mm to the right and left of the center of the gel, yielding similar values (∼40 kPa). There was not a statistical difference in opacity at those three points. Second, sparsely crosslinked alginate gels were exposed to a high concentration calcium chloride solution through a large (5 mm) circle stencil (**Figure 6B**). This yielded a radial gradient in opacity. The elastic modulus was calculated at the center of the gel and 1 and 3 mm to the right edge of the stencil. In the center of the gel, the elastic modulus was high (30 kPa), but further away the elastic modulus was much lower (∼10 kPa). These gels showed larger changes in opacity. Finally, sparsely crosslinked alginate gels were exposed to 60 mM NaCl (**Figure 6C**). This yielded no increase in opacity. The elastic modulus was calculated at the center of the gel and 3 mm to the right and left of the center of the gel, yielding similar values (∼1.5 kPa). All points both before and after addition of the 60 mM NaCl showed no visible difference in opacity. This demonstrates that elastic modulus differs spatially across gradients formed using stencils and that there is likely a relationship between opacity and elastic modulus.

**Figure 6:**
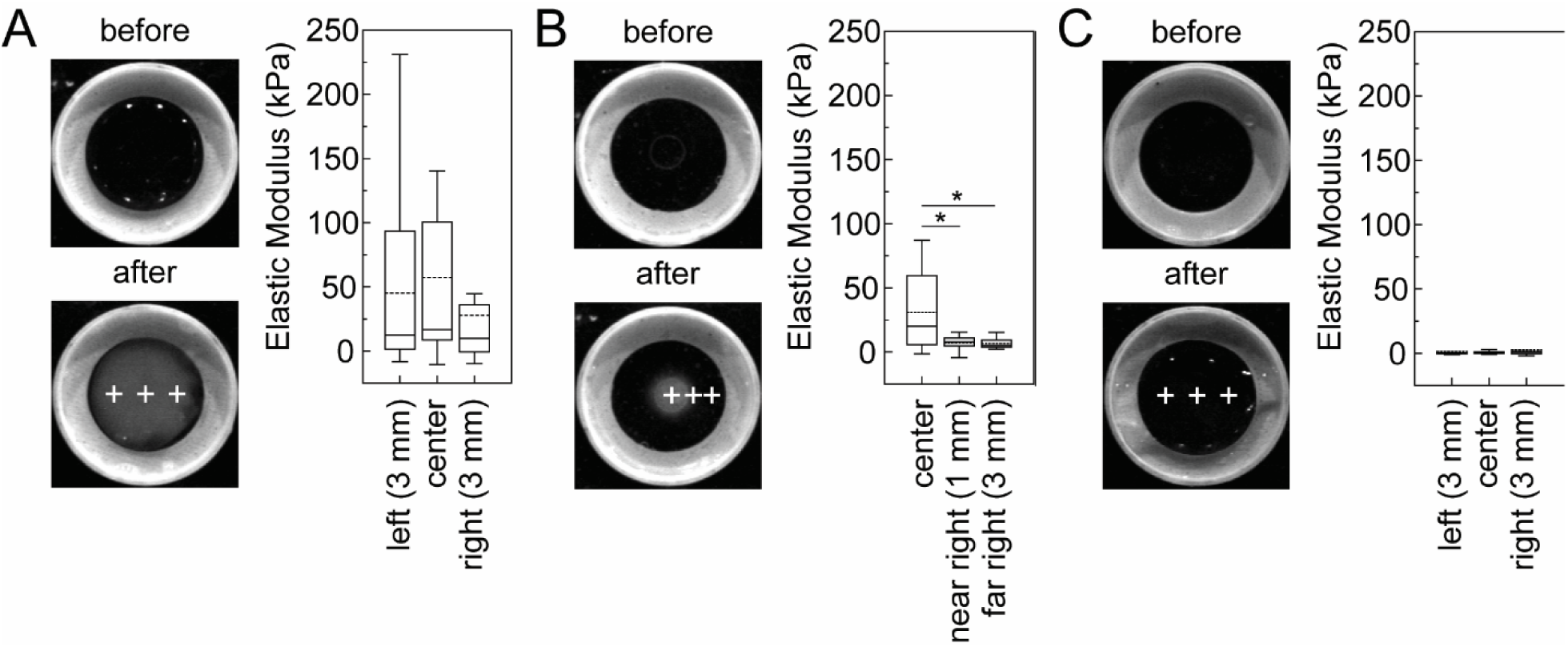
Stencils can create spatial mechanical property gradients in alginate gels. Imaging and mechanical property measurements were conducted on alginate gels (20 mg/ml alginate cross-linked with 20 mM CaSO_4_) that were A) exposed to a solution of 80 mM CaCl_2_ with no stencil, B) exposed to a solution of 80 mM CaCl_2_ above a 5 mm circular stencil or C) 60 mM NaCl for 3 h. Images were recorded before and after exposure to the solution. Mechanical properties were assessed at different positions denoted by the crosses in the image. Images are 25.4 mm by 25.4 mm. The 80 mM CaCl_2_ no stencil condition mechanical properties were determined from four days of experiments on four different gels probed at least three different times (*N_left_* = 12, *N_center_* = 13, *N_right_* = 13). The 80 mM CaCl_2_ stencil condition mechanical properties were determined from three days of experiments on three different gels probed at least three different times (*N_center_* = 14, *N_near right_* = 10, *N_far right_* = 9). The 60 mM NaCl no stencil condition mechanical properties were determined from two days of experiments on two different gels probed at least three different times (*N_left_* = 12, *N_center_* = 9, *N_right_* = 7). Box plots: Upper and lower error bars are minimum and maximum values, top and bottom of the box represent the 75^th^ and 25^th^ percentile, the line is the median and the dashed line is the mean. Conditions that resulted in *t*-tests with *p* < 0.05 are denoted with asterisks.

Given that both the elastic modulus and intensity decrease as a function of distance from the edge of the stencil, we overlaid elastic modulus and normalized intensity (**Figure 7A**). Similar decreases are observed for both opacity and elastic modulus. When the elastic modulus for each measurement is plotted against the corresponding normalized intensity, the result is a sigmoidal curve where the intensity increases with elastic modulus (**Figure 7B**). The half max for the sigmoidal fit is 8 kPa, indicating a steep increase in opacity. It appears that while opacity indicates stiffness changes, this range only spans from 5 kPa to 15 kPa.

**Figure 7:**
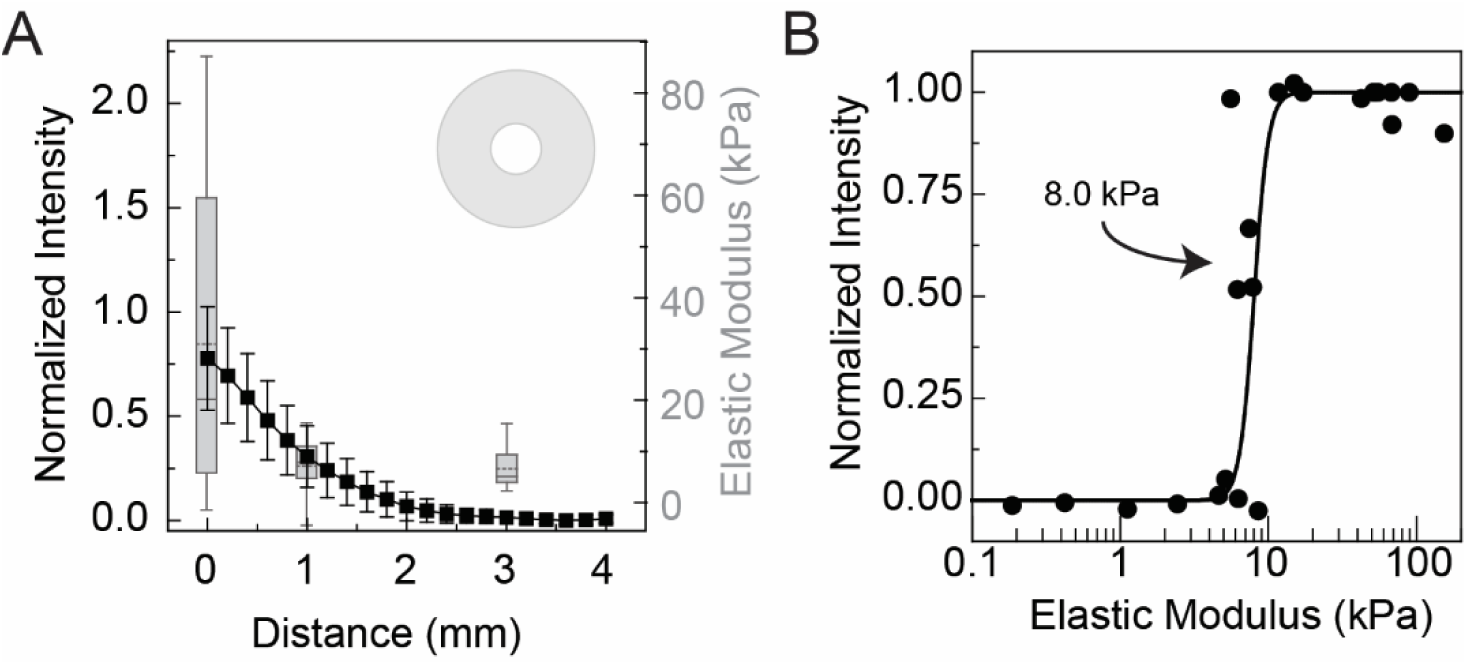
Opacity correlates with elastic modulus. A) Alginate gels (20 mg/ml alginate cross-linked with 20 mM CaSO_4_) were exposed to a solution of 80 mM CaCl_2_ with a 5 mm diameter circular stencil (top right) for 3 h and the opacity was measured (black squares, left) along with the elastic modulus (grey box plot, right) at different distances from the stencil edge. Error bars represent 95% confidence intervals. Box plots: Upper and lower error bars are minimum and maximum values, top and bottom of the box represent the 75^th^ and 25^th^ percentile, the line is the median and the dashed line is the mean. B) Opacity is plotted as a function of elastic modulus for positions in alginate gels (20 mg/ml alginate cross-linked with 20 mM CaSO_4_) that were exposed to a solution of 80 mM CaCl_2_ with no stencil, exposed to a solution of 80 mM CaCl_2_ above a 5 mm circular stencil or 60 mM NaCl for 3 h. The model is an empirical sigmoidal fit with the half max shown.

### 3.5 Cell invasion into the alginate gel is stiffness dependent

After establishing that gradients in mechanical properties can be generated using this stenciling approach, cellular responses to these gradients were investigated. Alginate is weakly cell adhesive so the alginate was functionalized with collagen and then weakly crosslinked using the CaSO_4_ solution. A 5 mm diameter stencil and 80 mM CaCl_2_ was used to imprint a radial gradient in the center of the alginate gel. Overlapping images across the diameter of the collagen functionalized alginate gel were taken of breast cancer cells (MDA-MB-231) on the surface of the gel and stitched together (**Figure 8A**). When alginate was not functionalized with collagen, there were fewer cells on the surface at 4 h compared to the collagen functionalized alginate gels (**Figure 8B**). Interestingly after 24 h the number of cells on the surface of the collagen gels significantly decreased. The decrease in cell number could either be due to cell death or cell invasion. Consequently, we fixed and stained cells for F-actin at 4 and 24 h and used confocal microscopy to image cells at different positions corresponding to different stiffnesses previously assessed by AFM (**Figure 8C**). Images were taken at different depths at these three spatial positions. Cells are seen in the center of collagen functionalized alginate gels at the surface at 4 h. However, they are no longer present at 24 h at the same position (**Figure 8D**). Conversely, cells are not seen in the center of the collagen functionalized alginate gel 150 μm under the surface at 4 h. However, they are observed at that depth at 24 h, indicating invasion. We quantified the cell density at different depths (**Figure 8F**). Very little cell attachment is shown in the absence of collagen. However, there is a robust decrease in cell density as a function of depth for all three positions at 4 h on collagen functionalized alginate gels. In the center, where the stiffness is higher, there is more robust invasion of cells into the collagen functionalized alginate gel resulting in both a decrease in cell number at the surface and an increase in cell number at lower depths at 24 h. There is approximately similar invasion at 1 mm from the stencil edge, however, more cells remain closer to the surface. Interestingly, there is very little cell invasion at 3 mm from the stencil edge, resulting in a depth dependent cell density that is roughly similar between 4 and 24 h. This demonstrates that imprinting of mechanical property gradients using this unique approach can create spatially dependent invasion patterns in breast cancer cells.

**Figure 8:**
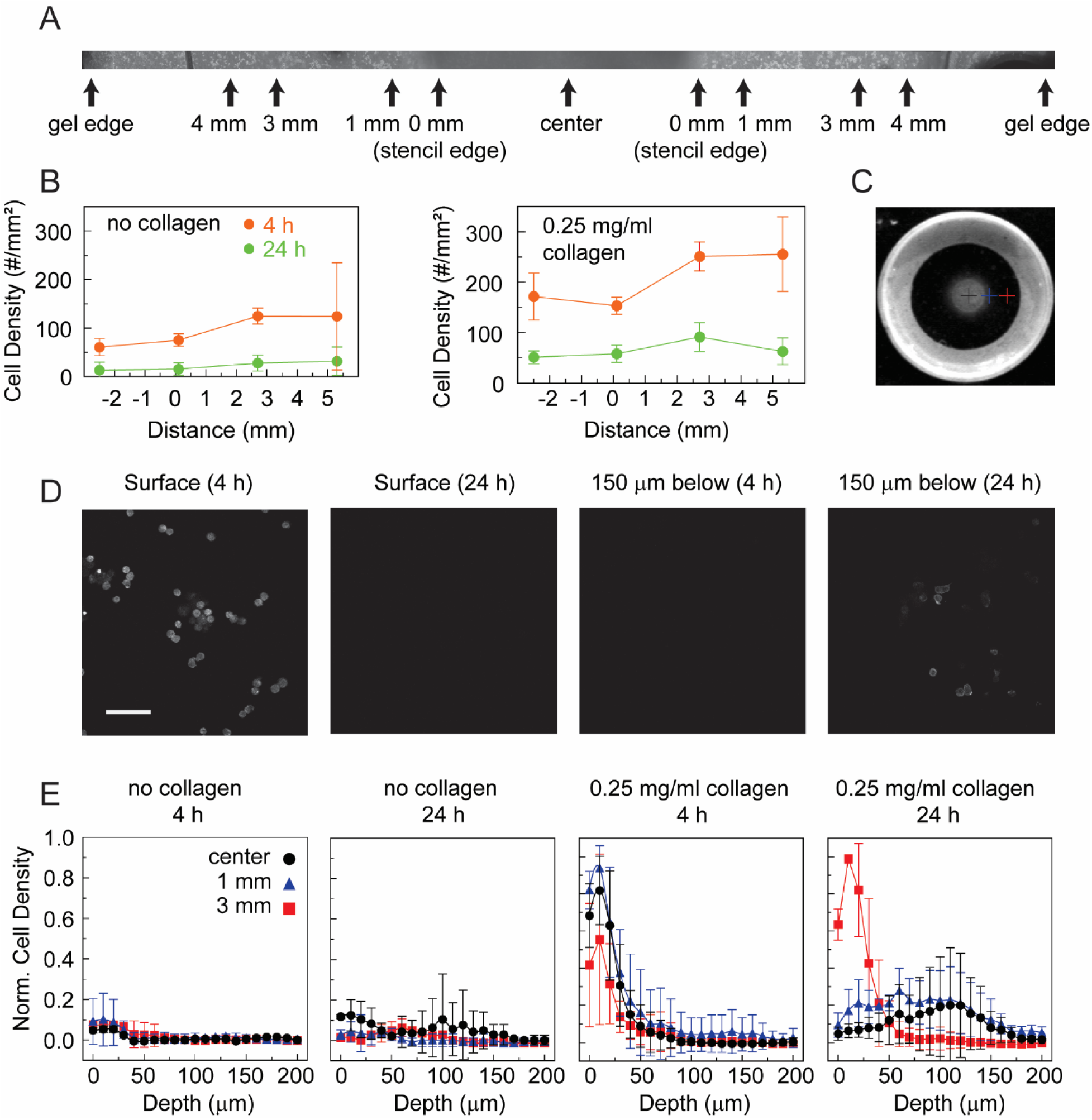
Stiffness gradients create a gradient in cancer cell invasion efficiency. Breast cancer cells (MDA-MB-231) were plated on the surface of collagen functionalized alginate gels with spatial stiffness gradients. A) Phase contrast images were taken and stitched across the entire gel. Positions where measurements were taken and reported are marked. B) Cell attachment density was quantified at different positions by examining the stitched images of alginate gels with and without collagen functionalization at both early (4 h) and late (24 h) timepoints. (no collagen: *N_days_* = 2, *N_gels_* = 3, *N_points_* = 6, collagen: *N_days_* = 2, *N_gels_* = 4, *N_points_* = 8) C) Image of the alginate gel is marked at different positions denoted by the crosses to indicate where invasion was assessed. Images are 25.4 mm by 25.4 mm. D) Confocal fluorescence images were taken of cells fixed and stained with phalloidin on alginate gels functionalized with collagen in the center of the gel at different times and at different depths. The scale bar is 100 μm. E) Normalized cell density was quantified for alginate gels with and without collagen functionalization measured at the center of the gel (black circles), 1 mm from the stencil edge (blue triangles) and 3 mm from the stencil edge (red squares) at different depths (no collagen: *N_days_* = 2, *N_gels_* = 2, *N_points_* = 2, collagen: *N_days_* = 2, *N_gels_* = 3, *N_points_* = 4). Error bars are 95% confidence intervals.

## 4. Discussion

In this paper we created an approach to generate tunable and reproducible gradients in both thick and thin alginate gels using an easy stenciling approach. We characterized the crosslinking kinetics a number of different ways and showed that geometry and gradient development time can tune both the average opacity and opacity gradient steepness in known and reproducible ways. We measured Young’s modulus in both bulk materials as well as spatially, demonstrating that the opacity gradient mapped to the stiffness gradient and quantified the range of stiffness over which opacity was a good proxy for stiffness. We demonstrate a method of functionalizing stiffness gradient alginate with collagen resulting in spatially dependent cell invasion into the alginate gel.

Opacity changes in alginate have been observed previously. However, the mechanism for changes in opacity are obscure. Opacity changes are likely due to increases in light scattering caused by local contraction and densification of the alginate network. This is likely, given that hydrogel opacity varies with polymer molecular weight, concentration and crosslinking density and high water content hydrogels have refractive indices similar to that of water[35]. Gel contraction is likely driven by minimizing free energy due to the decrease in entropy associated with the crosslinks[36]. Interestingly, we observed flow from outer edges in areas that were not opaque towards the opaque center when using radial stencils. This flow pattern changes based on the shape of the stencil. This directed contraction could be leveraged as a way to locally stimulate cells by inducing shear or compressive stresses upon the cells. In addition, interpenetrating networks composed of alginate and fiber forming polymers like collagen or fibrin could be assembled. The spatially controlled contraction that occurs within these alginate gels could be used to align or structure the fiber network in these composite networks. Consequently, the contraction may be leveraged in interesting ways. Applications that require either cell embedding or assembly of the gradient on tissue will need to account for this contraction and if undesired, approaches to mitigate it by using a stiffer initial gel or altering ionic strength of the crosslinking gel could be used.

As a demonstration of the utility of these gradients we examined cancer cell invasion into these alginate gradient gels. We showed spatial control over invasion through the stiffness gradient formation. Cancer cells in regions that were stiffer invaded more efficiently than in regions that were softer. This broadly agrees with what has been seen in other systems examining the role of stiffness in controlling cell migration and invasion[37]. Others have shown invasion into alginate gels that is stiffness dependent although under certain circumstances higher stiffness either stunts migration or facilitates it [38–40]. This seems to be dependent on the extracellular matrix additions to the alginate including fibrillar collagen or decellularized extracellular matrix. More recently, the plastic nature of the extracellular matrix has become an appreciated mechanical modulator of cancer cell migration to complement the already established role for elastic behavior in driving cancer invasion[41]. Future efforts focusing on controlling viscoelastic properties spatially using this technique will likely uncover novel roles for the spatial regulation of cell behavior through plastic deformations of the extracellular matrix.

One potential use for these gradient gels would be in experiments exploring fundamental aspects of durotaxis. While there are many examples of creating 2D stiffness gradients to explore how signaling or the cytoskeleton regulates durotaxis, there are many fewer examples of mechanistic studies of durotaxis in 3D. The Young’s modulus probed in this work (0.5-50 kPa) and in particular the Young’s modulus over which opacity is a good proxy for stiffness (5-15 kPa) matches well with other work characterizing Young’s modulus in alginate gels[42–44] and overlaps nicely with soft tissues and the range of mechanosensitivity of many types of cells[45]. Alginate is a relatively inert molecule and many cells do not attach to it. However, by functionalizing the alginate gel with collagen, cells can invade the gel, allowing for studies of durotaxis in 3D. Consequently, there is an opportunity to attach extracellular matrix molecules to the alginate to study durotaxis across a variety of ligand-receptor pairs. Furthermore, there are opportunities to assemble interpenetrating networks composed of both alginate and collagen and create similar gradients, assessing whether cells can sense stiffness gradients in alginate through the collagen network[44,46]. After stiffness gradient imprinting, different stencils could be used to generate similar or spatially distinct gradients of soluble factor to explore how cells integrate durotactic and chemotactic cues. Collagen fibers within the networks could be aligned using magnetic fields[47,48] or local mechanical rotation[49], allowing to probe competition or cooperation between contact guidance and durotaxis. Understanding how different cells integrate these signals and leverage overlapping or distinct intracellular signaling networks is an understudied area and is absolutely critical for the assembly of complex tissues.

Another potential use for this system would be as an advanced wound dressing. There are other examples of systems that have used soluble gradients to direct cell migration in the hope of enhancing wound healing[50,51]. However, to our knowledge there are no reports of efforts using stiffness gradients to enhance wound healing, even though stiffness gradients are features of healing wounds[14]. As mentioned above, alginate has already been used as a wound dressing, but mainly for its ability to absorb exudate. Either attaching extracellular matrix or assembling an interpenetrating network would allow for its use as a provisional matrix, guiding cell movement into the provisional matrix through gradients of stiffness. The versatility of this technique could be leveraged to generate gradients that best match the geometry and mechanics of the wound. The ease in creating these gradients will allow non-experts the ability to create made-to-order wound dressings.

## 5. Conclusions

Stencils can be used to generate gradients in opacity within alginate gels. These stencils can be easily designed and fabricated using a common craft cutter. Different stencil shapes result in different gradients in opacity that can be imprinted into both thick and thin alginate gels of arbitrary shape. The steepness of the opacity gradient as well as the maximum opacity can be controlled based on reproducible crosslinking kinetics regulated through calcium concentration and gradient developing time. Calcium crosslinking results in both opacity changes as well as increases in elastic modulus in the bulk hydrogel. Spatial gradients in elastic modulus can also be imprinted into alginate gels using this stenciling approach. Opacity correlates with elastic modulus, allowing it to be used as a proxy for local elastic modulus. Finally, gradient alginate gels can be functionalized with collagen facilitating cancer cell invasion that occurs at different rates depending on the spatial position of the cell.

## Acknowledgements

We acknowledge Megan Wolfe, Sarah Fangrow, Maria Lebedeva, Madison Karamagianis, Flora Kafunda, Clarissa Carolan, Claire Allen from the Freshman Honors Research Program. We also acknowledge Margie Carter and the Office of Biotechnology and the Roy J. Carver High Resolution Microscopy Facility for help with the confocal microscopy.

## Abbreviations

ECM: extracellular matrix
UV: ultraviolet
VIS: visible
PBS: phosphate buffered saline
DMEM: Dulbecco’s modified Eagle medium
FBS: fetal bovine serum
APTES: aminoproplytriethoxysilane
NHS: N-hydroxysuccinimide
EDC: 1-ethyl-3-(3-dimethylaminopropyl) carbodiimide
FTIR: Fourier transform infrared
ATR: Attenuated total reflectance
4NB: 4-nitrobenzaldeyhde
AFM: Atomic force microscopy
MES: 2-(*N*-morpholino) ethanesulfonic acid
EGTA: Ethylene Glycol Tetraacetic Acid
TBS: Tris Buffered Saline
STED: Stimulated Emission Depletion

## Data Availability

## Author Contributions

Z.O.: Conceived, conducted, analyzed the experiments and interpreted the data across figures 1, 3, 6-8, S1 and S5-S8, wrote the article, prepared the figures and edited the article. T.P.: Conceived, conducted, analyzed the experiments and interpreted the data across figures 4, 5 and S4, wrote the article, prepared the figures and edited the article. J.Z.: Conceived, conducted, analyzed the experiments and interpreted the data across figures 6 and S8, wrote the article and edited the article. A.F.: Conceived and conducted experiments and interpreted the data in figure 8, wrote the article and edited the article. F.R.N.: Conceived and conducted experiments in figure 8, wrote the article and edited the article. T.K.: Conceived the experiments in figures 1 and 3 and edited the article. N.J.: Conceived, conducted, analyzed the experiments and interpreted the data in figure 2. T.G.: Conceived, conducted, analyzed the experiments and interpreted the data in figure S3. M.T.: Conceived, conducted, analyzed the experiments and interpreted the data in figure S2, wrote the article and edited the article. E.M.: Conceived and printed molds used across all figures, wrote the article and edited the article. L.M.: Conducted the experiments in Figure S6. A.K.: Conducted the experiments in Figure S6. A.M.: Conceived molds used across all figures and edited the article. J.R.: Conceived and interpreted the data across figures 6 and S8. I.C.S.: Conceived, analyzed the data and interpreted the data across all figures, wrote the article, prepared the figures and edited the article.

---

**Supplementary Figure 1:**
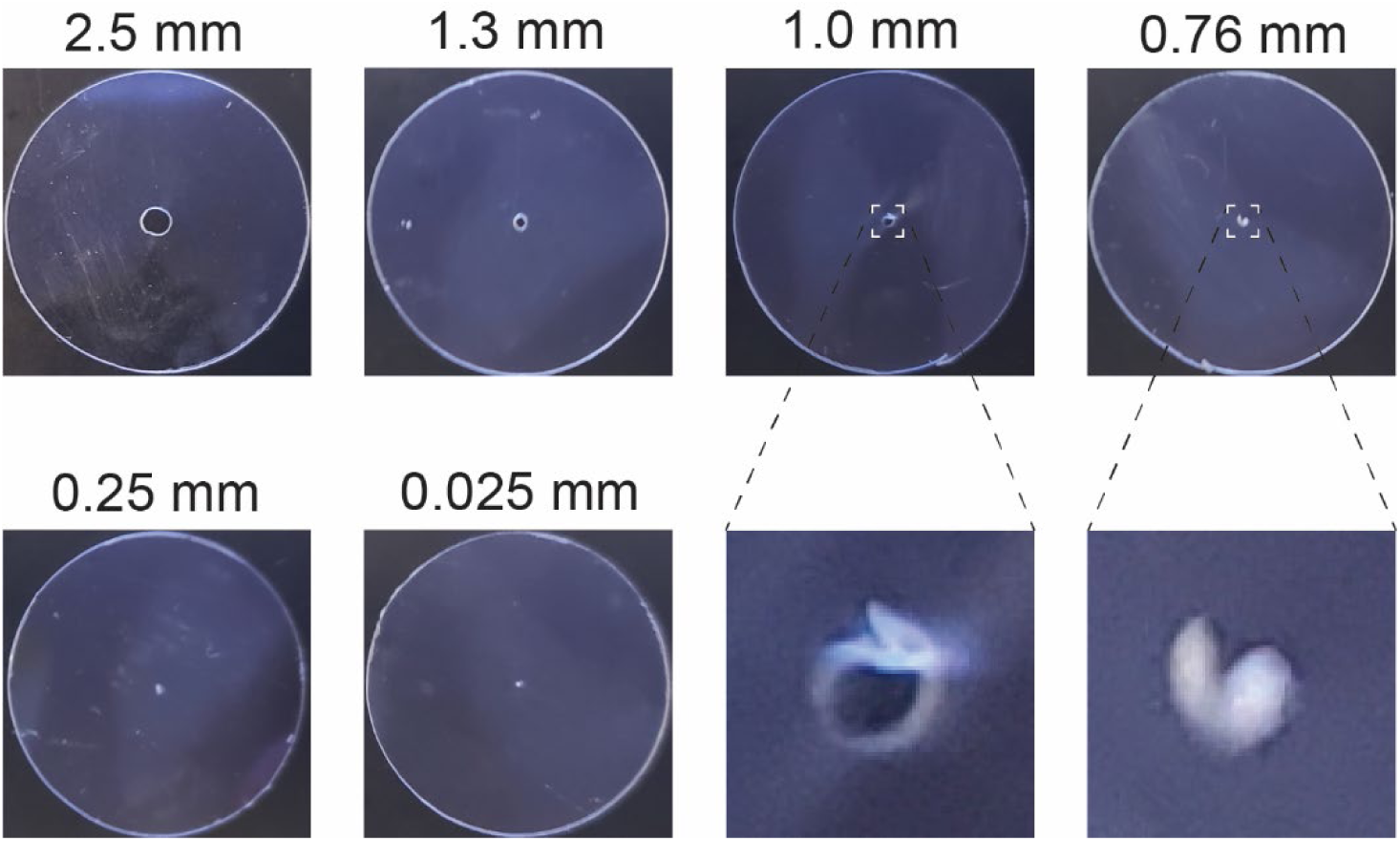
Craft cutter stencil cutting resolution. Circles of various diameters were cut into the stencil material. Enlarged regions of selected stencils are shown. Images are 25.4 mm by 25.4 mm.

**Supplementary Figure 2:**
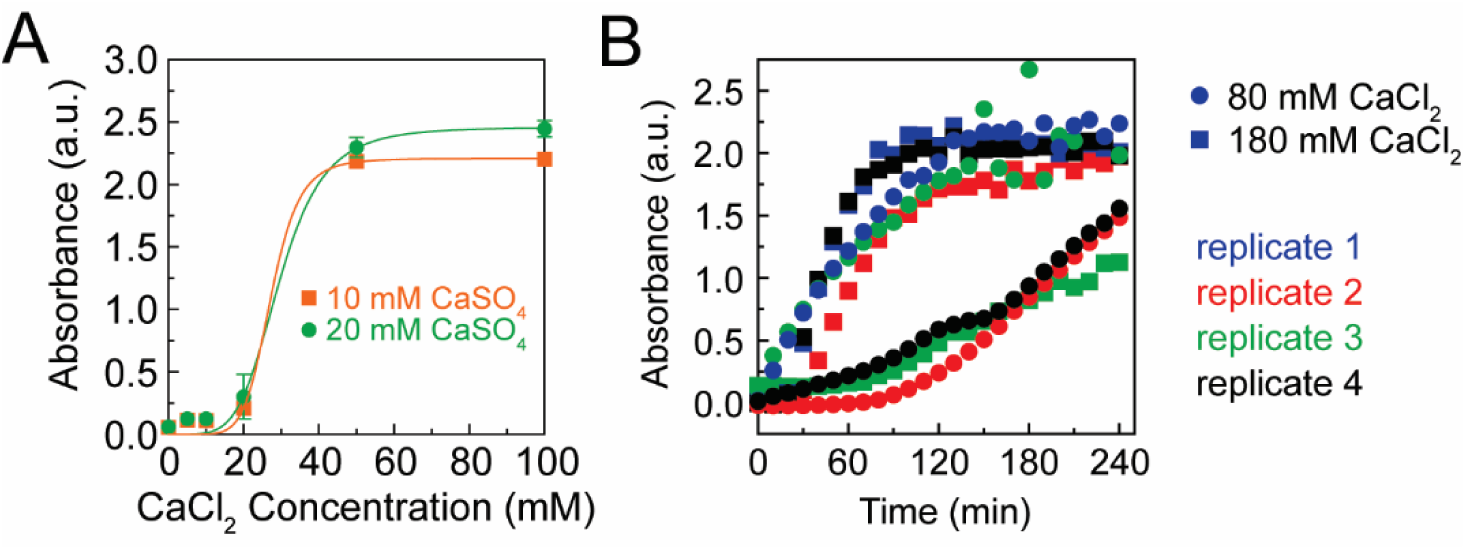
Measuring kinetics of alginate polymerization using absorbance. A) Dose response of the absorbance at 750 nm of an alginate gel (20 mg/ml alginate) sparsely cross-linked with 20 mM CaSO_4_ (green) or 10 mM CaSO_4_ (orange) as a function of exposure to different concentrations of CaCl_2_ after 16-24 h (*N_gels_* = 3-5). Error bars represent 95% confidence intervals. B) Absorbance at 750 nm of alginate gels (20 mg/ml alginate cross-linked with 20 mM CaSO_4_) after exposure to 80 or 180 mM CaCl_2_ as a function of time.

**Supplementary Figure 3:**
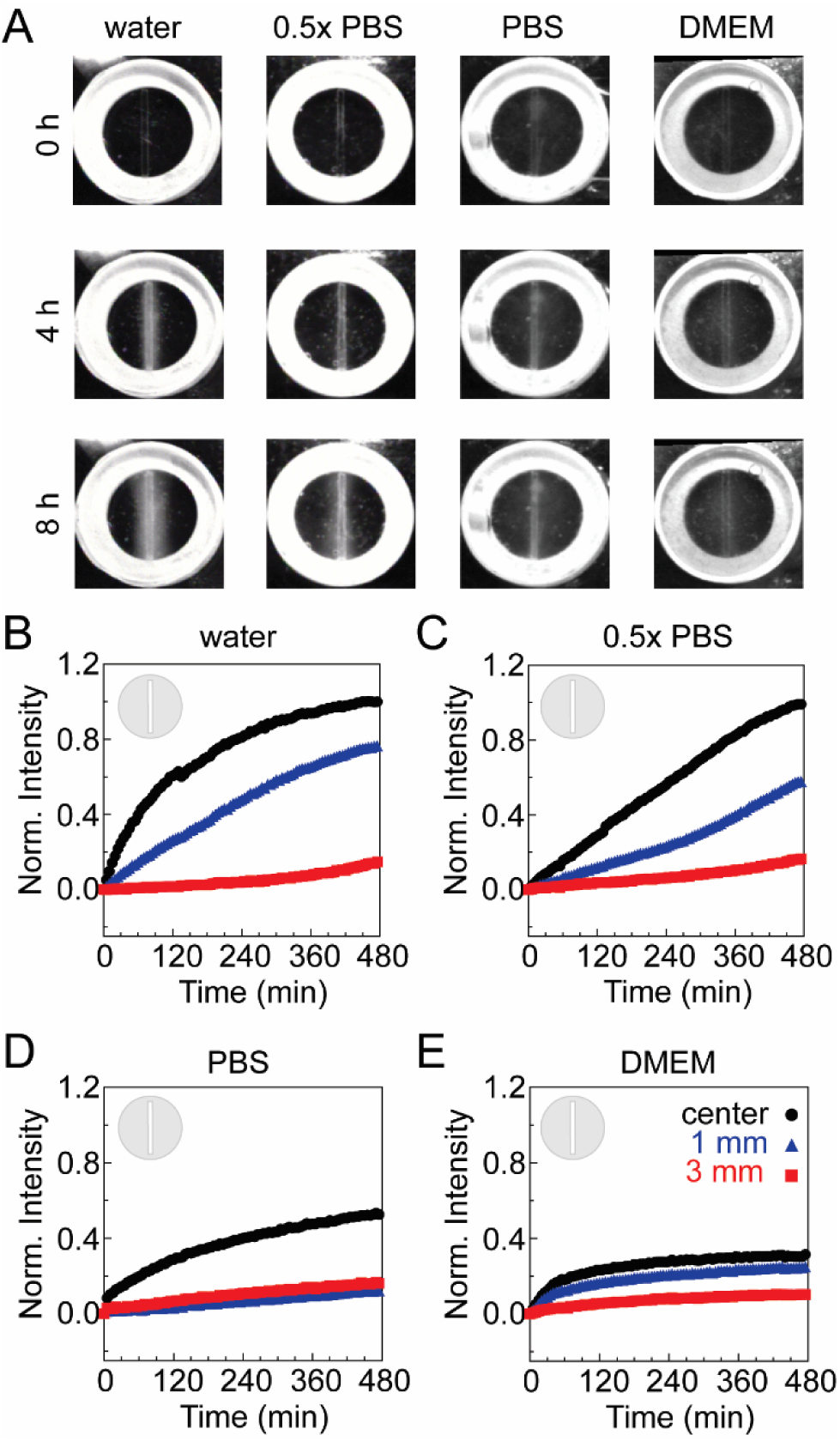
Gradients in opacity produced in other buffers. A) Images of alginate gels (20 mg/ml alginate cross-linked with 20 mM CaSO_4_) in different buffers (water, 0.5X PBS, PBS and DMEM) at various times (0, 4 and 8 h) after exposure to an 80 mM CaCl_2_ solution in different buffers (water, water, PBS and DMEM) above the stencil. Images are 25.4 mm by 25.4 mm. Normalized intensity over time of gels in B) water, C) 0.5x PBS, D) PBS and E) DMEM with a line stencil on the alginate gels at distances of 0 (center), 1 and 3 mm from the center of the gel.

**Supplementary Figure 4:**
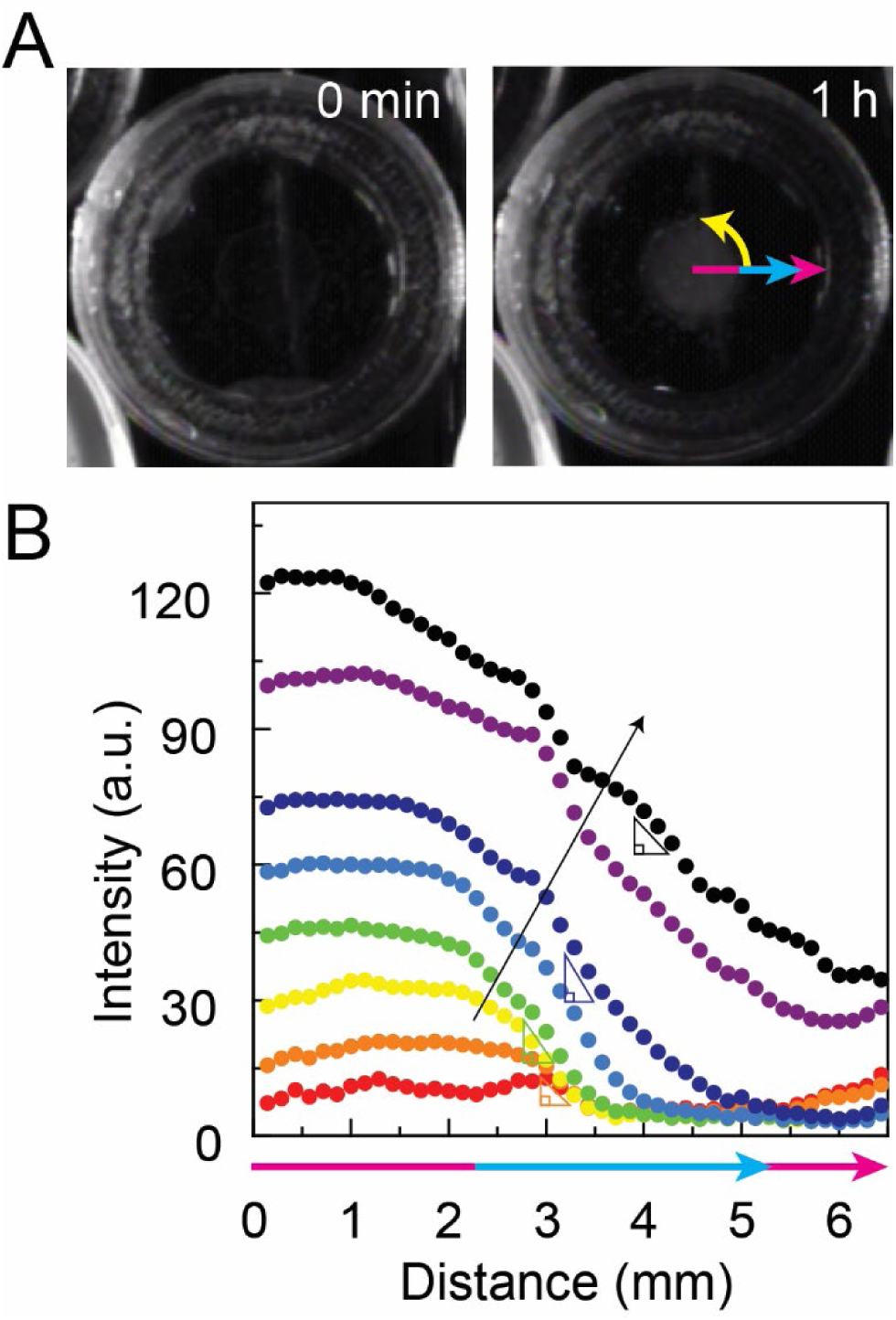
Description of analysis approach to quantify the opacity radial profile. A) Representative images of alginate gels (20 mg/ml alginate cross-linked with 20 mM CaSO_4_) before (0 h) and after (1 h) exposure to an 80 mM CaCl_2_ solution above the stencil. The entire radial linescan (magenta arrow), the radial linescan used for data analysis (blue arrow) and the direction of angle sweep to average the radial opacity profile over all angles. Images are 25.4 mm by 25.4 mm. B) Normalized intensity averaged over all angles as a function of distance at various times ranging from short times (0 h, red) to long times (16 h, black). Triangles represent where an absolute gradient is calculated for a specific position.

**Supplementary Figure 5:**
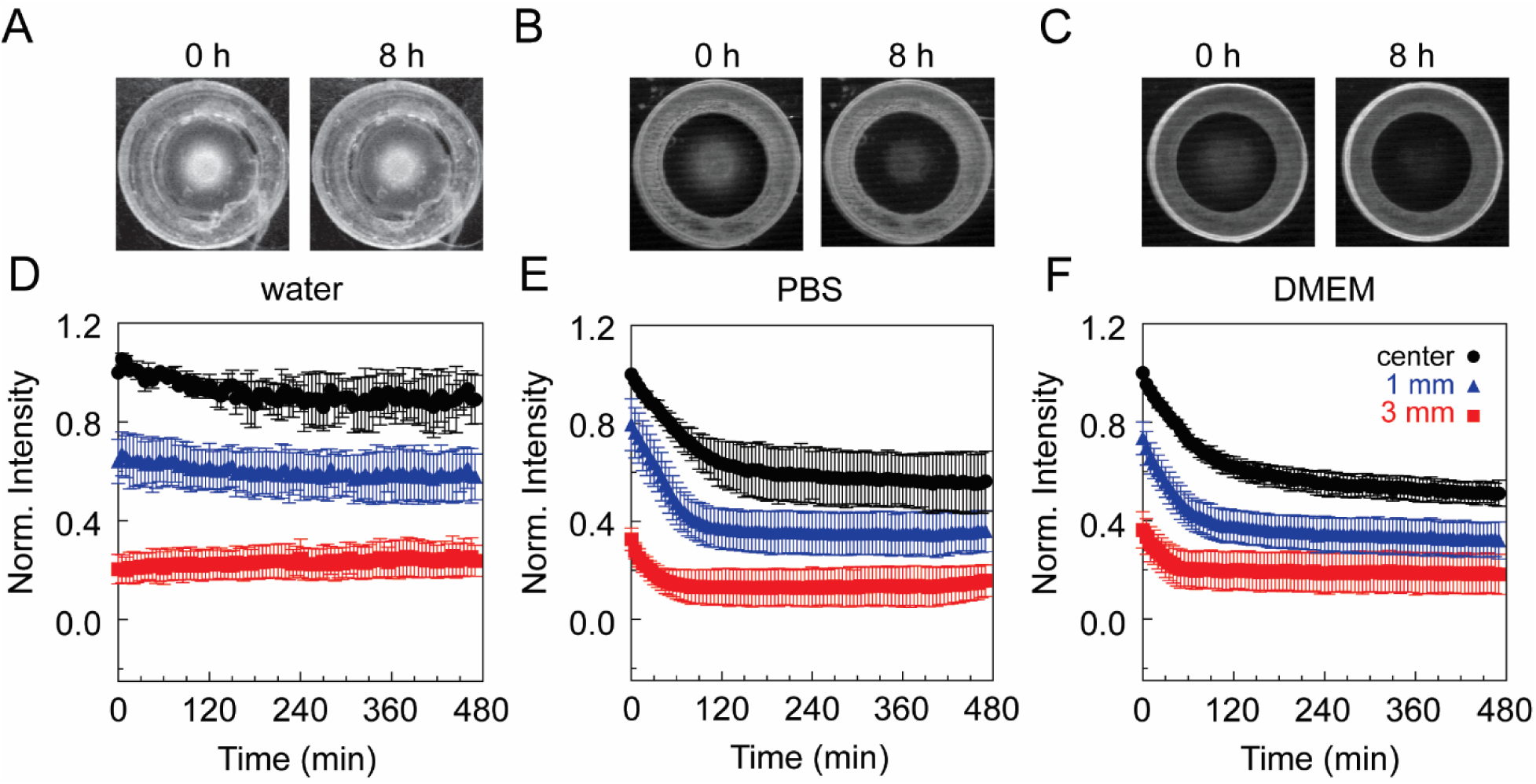
Measuring opacity changes after removal of calcium from solution above gels. Representative images of alginate gels (20 mg/ml alginate cross-linked with 20 mM CaSO_4_) after 8 h exposure to an 80 mM CaCl_2_ solution above the stencil (marked as 0 h) and subsequent exposure to A) water, B) PBS or C) DMEM (marked as 8 h). Images are 25.4 mm by 25.4 mm. Normalized intensity over time at distances of 0 (center), 1 and 3 mm from the center of the gel in D) water (*N_days_* = 2, *N_gels_* = 3, *N_points, center_* = 3, N*_points, 1mm or 3mm_* = 6), E) PBS (*N_days_* = 2, *N_gels_* = 3, *N_points, center_* = 3, N*_points, 1mm or 3mm_* = 6) and F) DMEM (*N_days_* = 2, *N_gels_* = 4, *N_points, center_* = 4, N*_points, 1mm or 3mm_* = 8). Error bars represent 95% confidence intervals.

**Supplementary Figure 6:**
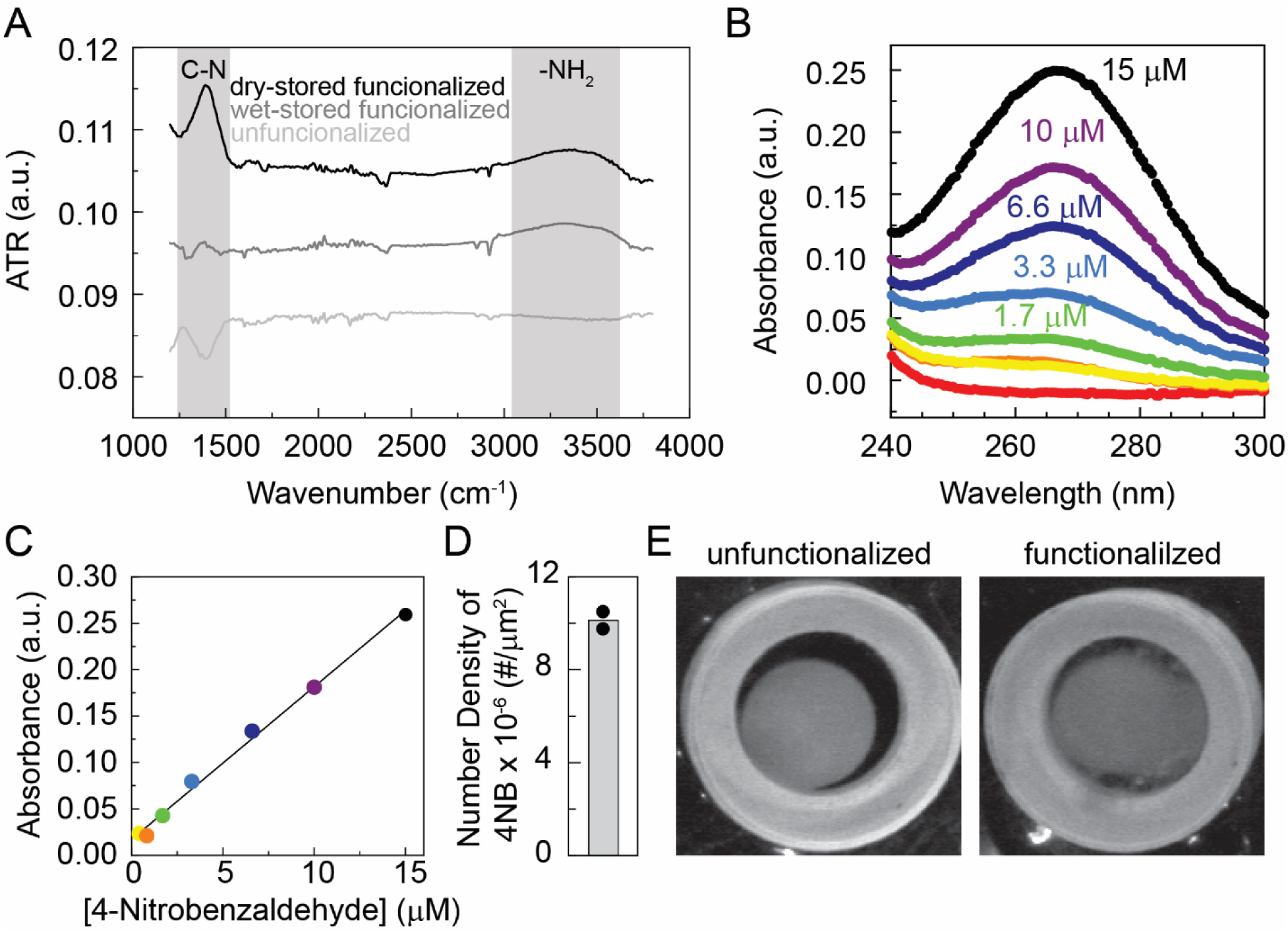
Characterization of functional attachment of alginate to glass. A) ATR signal from FTIR spectroscopy of functionalized borosilicate glass kept dry or wet and unfunctionalized glass. Probable functional groups that cause the largest peaks in the ATR are denoted above. B) Absorbance scan of different concentrations of 4-nitrobenzaldehyde (red is 0 μM). C) Calibration curve generated by measuring absorbance at 270 nm as a function of 4NB concentration with a linear fit. D) Quantification of number density of amine groups as calculated by the amount of 4-nitrobenzaldehyde (4NB) hydrolyzed from the surface. The black dots represent two independent experiments and the grey bar is the mean. E) Representative images of alginate gels (20 mg/ml alginate cross-linked with 20 mM CaSO_4_) after 8 h exposure to an 80 mM CaCl_2_ solution on either unfunctionalized glass or functionalized glass. Images are 25.4 mm by 25.4 mm.

**Supplementary Figure 7:**
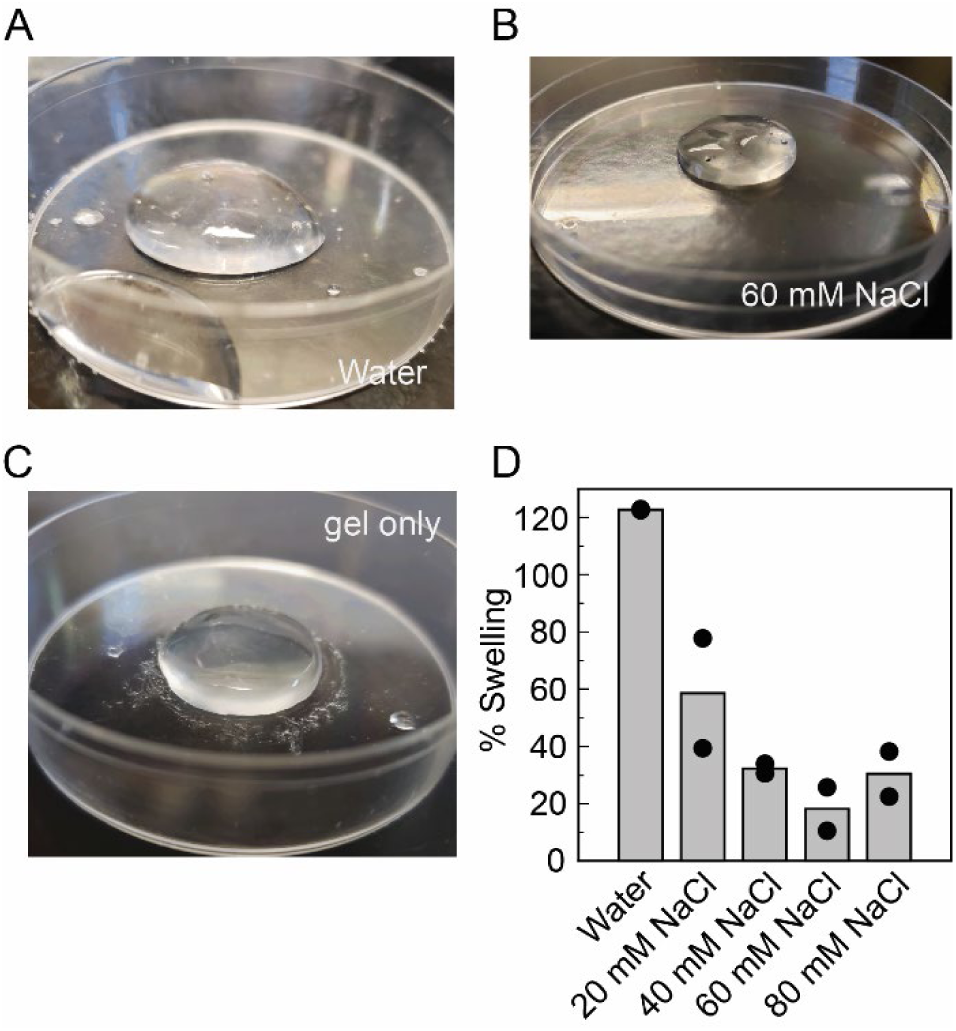
Swelling of alginate gels in different buffers. Representative images for alginate gels (20 mg/ml alginate crosslinked with 20 mM CaSO_4_) made in a 60 mm dish A) after exposure to water for 12 h, B) after exposure to 60 mM NaCl for 12h and C) initially at time 0 h. D) The average percent swelling (grey bars) and individual sample percent swelling (black circles) calculated at the mass of additional water in the gel divided by the initial mass of the gel x 100%.

**Supplementary Figure 8:**
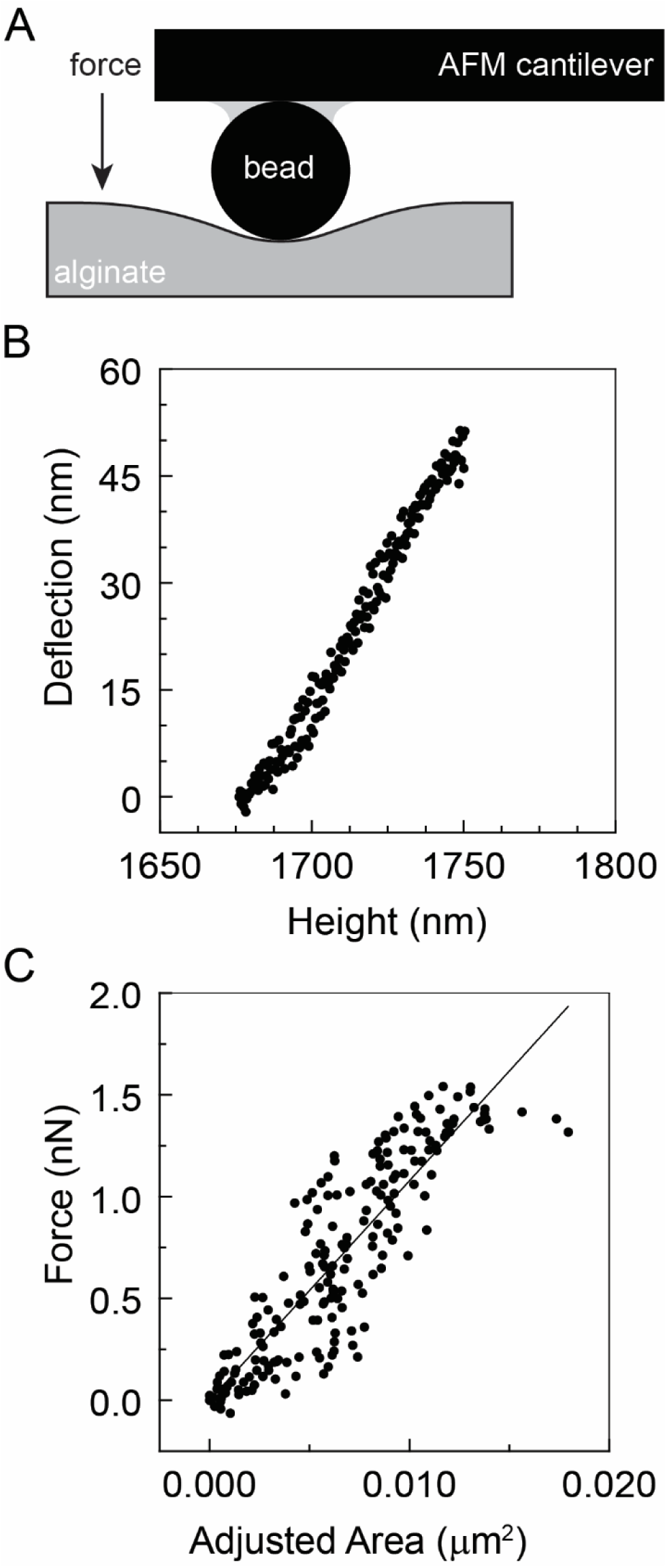
Measurement of elastic modulus using atomic force microscopy. A) Schematic showing the deformation experiment using AFM. B) Sample graph of cantilever deflection as a function of height. C) Sample graph of force as a function of adjusted area, where the slope is the Young’s modulus of the material.

